# Antennal sensilla diversity in diurnal and nocturnal fireflies (Coleoptera, Lampyridae)

**DOI:** 10.1101/2024.05.12.593785

**Authors:** Yelena M. Pacheco, Ethan Mann, Luiz F. L. Da Silveira, Seth M. Bybee, Marc A. Branham, Joseph V. McHugh, Kathrin F. Stanger-Hall

**Author notes:** Corresponding author (YMP), ORCID ID 0000-0003-4552-0715.

## Abstract

Insects use their antennae to collect environmental information. While the structural diversity of insect antennae is immediately obvious, the diversity of the minute antennal sensilla that interact with the environmental stimuli and translate them into sensory input, is largely unknown for many insect groups. This includes the beetle family Lampyridae, which includes nocturnal species that use bioluminescent signals during mate search, and diurnal species that rely exclusively on pheromones to identify and locate a potential mate. Diurnal species tend to have relatively larger antennae, and diurnal males have larger antennae than their females. It is generally assumed that antennal size reflects sensilla numbers, but this remains to be tested. Here we use Scanning Electron Microscopy to document the sensilla diversity of both males and females of three diurnal and four nocturnal firefly species, as well as total sensilla numbers, densities and their distribution along the antenna. We identified 14 sensilla morphotypes across the seven species, including 12 morphotypes that are new for Lampyridae. Mechanosensilla (3 morphotypes) were the most abundant and conserved sensilla across firefly species, and the distribution of chemosensilla (9 morphotypes) was unexpectedly variable across species. We hypothesized that the differences in mating signals between diurnal and nocturnal fireflies would be reflected in their chemosensilla counts or densities. As predicted, diurnal and nocturnal fireflies did not differ in their mechanoreceptor counts or densities, nor did males and females. In contrast, firefly males had significantly more chemoreceptors (and higher densities) than females and the interaction term (activity by sex) was also significant: diurnal males had significantly more chemoreceptors than nocturnal males, highlighting the importance of pheromones for diurnal species. Based on a series of predictions, we also identified a pheromone sensilla candidate for each species that will facilitate functional testing in future studies.

## Introduction

Antennae are major sensory organs of insects and are remarkably diverse in both form and function. Antennae are composed of multiple segments (antennomeres) whose numbers range from 2-62 across beetle groups [1,2]. There is morphological and functional variation between individual antennomeres: the basal two antennomeres (scape and pedicel) contain muscle attachments that are used to support and move the antennae, and the antennomeres that make up the remainder of the antenna (the flagellomeres) are involved with sensing the environment. The flagellomeres can be highly modified [3,4], creating diverse antennal shapes across insect groups [5]. While the diversity of antennal types showcases different ways of increasing antennal surface to collect environmental information, the first point of contact for the different environmental stimuli are the sensilla, microscopic sensory structures on the antennal surface. Sensilla are extensions of the insect cuticle [6] and each sensillum represents a specialized accessory structure that translates specific environmental stimuli for specific sensory neurons [7]. Functional sensilla types include mechanoreceptors (pressure, touch), chemoreceptors (volatile and/or contact chemicals), thermoreceptors (temperature) and hygroreceptors (humidity and air pressure) [6–8].

The antennal sensilla of insects are diverse in both form and function. The basic sensilla anatomy includes a stalk, which can emerge directly from the antennal surface or from an elevated base. The stalk can vary in shape and length, in the presence or absence of grooves along the length of the stalk and may lack or show pores [6]. These characteristics, along with the cell morphology of their sensory neurons, are traditionally used to classify sensilla into broad morphological groups. In his seminal review, Schneider [6] described nine morphological groups of sensilla found in all insects. Some of these morphotypes are typically associated with specific functions across insects. For example, chaetica and campaniform sensilla tend to be primarily mechanoreceptors [6, 9, 10], coeloconica and capitular sensilla tend to be thermo- or hygroreceptors [6, 11–13], and trichodea, basiconica, and placodea sensilla are typically chemoreceptors [6]. Additional sensilla morphotypes have been described for different insect groups, either as new variants of described insect morphotypes, or as entirely new sensilla types. For example, a wide variety of sensilla morphotypes have been identified within beetles (25 in Cerambycidae [14], 16 in *Agriotes* Elateridae [15], 16 in Scarabaeidae [16] with sensilla chaetica, basiconica, and trichodea representing the most common sensilla types across these beetles. However, sensilla morphotypes greatly varied between families and even between species.

The specific function of individual sensilla morphotypes has been studied in relatively few beetle species. Sensilla chaetica are well established as mechanoreceptors [6, 11] and specific examples in Coleoptera include *Oryzaephilus surinamensis* (Silvanidae) and *Limonius aeruginosus* (Elateridae [17, 18]). Among chemoreceptors most functional studies have focused on the detection of plant volatiles, sex pheromones, and/or aggregation pheromones [19]. For example, sensilla trichodea detected sex pheromones in the pine weevil *Hylobius abietu* (Curculionidae [20]), and sensilla basiconica were shown to detect sex pheromones in the firefly *Photinus corruscus* [21]. In addition, sensilla coeloconica have been identified as thermoreceptors in *Siagona europaea* (Carabidae [22]), and as hygroreceptors in the firefly *Luciola cruciata* [23].

### Increasing antennal sensitivity for enhanced stimulus detection

Given the importance of antennae and their sensilla for insects to sense their environment, strong natural or sexual selection for the improved detection of relevant stimuli is expected [24]. One way to improve stimulus detection is by using more sensors. This could be achieved by increasing the surface area of the antennae [25, 26], thus increasing the number of sensilla and their associated sensory neurons on the antennal surface [27], while at the same time increasing the air space that the antennae can sample. The surface area of the antennae can be increased in one of three ways: (1) Increased number of antennal segments, (2) increased length of individual segments, or (3) the addition of side-branches to segments [2, 6]. In addition, stimulus detection could be further improved by (4) increasing the density of the sensilla on a given antennal surface area [25, 28] and/or by (5) increasing the size of the individual sensilla to maximize interaction with environmental stimuli and thus their sensitivity at threshold levels [10].

### Focus on Fireflies

We focus here on the antennal sensilla diversity of adult Lampyridae. Firefly species occur worldwide (except Antarctica) and exhibit a striking range of antennal diversity [2, 29]. This diversity includes both antennal shape and antennomere number. Within Coleoptera, the typical number of antennomeres is eleven [30], and indeed most beetle families have this fixed number across all species [1]. Among fireflies the number ranges from 7-62 antennomeres, with eleven being the most common (Fig 1; [2]). Within-species variation is known to occur in several firefly genera, including *Alecton*, *Microphotus*, *Pleotomus* and genera of Amydetinae [2]. To date, antennal sensilla diversity in fireflies has only been studied in the bioluminescent males of one firefly species, *Luciola cruciata* [23] from Asia. Iwasaki et al. [23] described seven sensilla morphotypes, including four mechanoreceptors, two chemoreceptors and one hygroreceptor. It is unknown whether the morphotypes of *L. cruciata* are representative of other firefly species and whether there are differences in sensilla types, numbers, or densities between males and females and/or between bioluminescent (nocturnal) and non-bioluminescent (diurnal) firefly species. Filling this gap is important, because the absence or presence of adult bioluminescence has implications for how firefly antennae are used.

**Fig 1.**
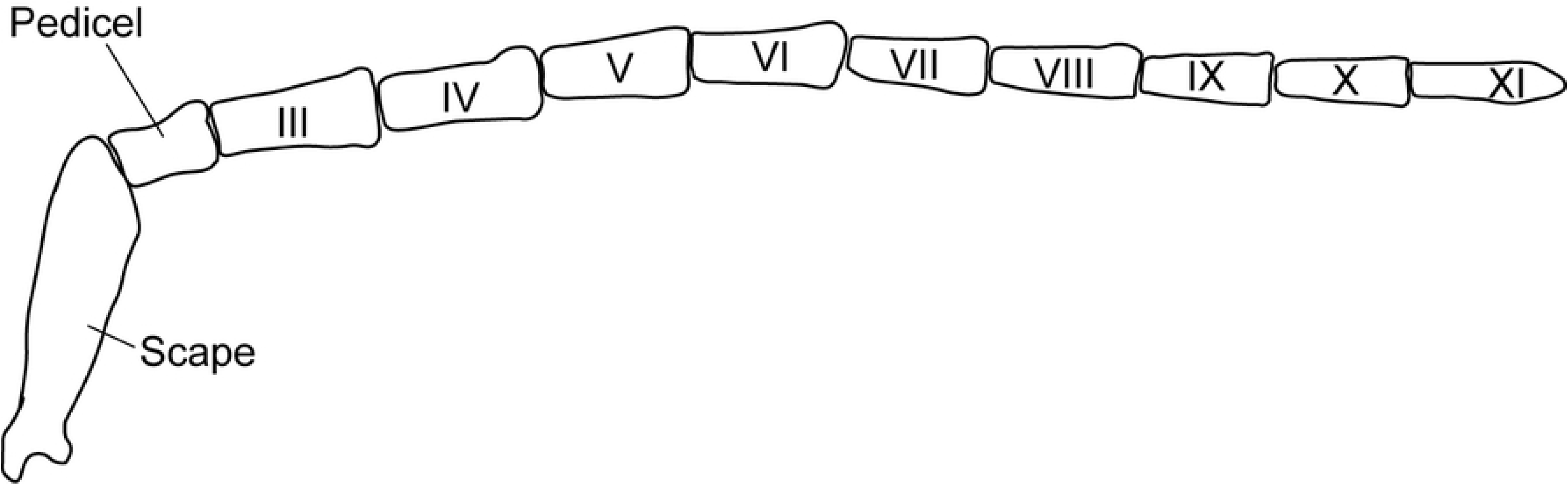
Diagram of adult firefly antennae with 11 antennomeres. Scape and pedicel represent the first two antennomeres.

Nocturnal firefly species are active at dusk or at night and are bioluminescent. They use prolonged glows [31] or species-specific flash patterns [32–34] as visual signals to attract and recognize conspecific mates. In contrast, most diurnal firefly species have non-bioluminescent adults that rely exclusively on pheromones to identify and locate a conspecific mate. In both signaling systems males actively look for females during mate search, while females are sedentary, and correlated with these behaviors, males have significantly larger eyes than their conspecific females [35]. Furthermore, bioluminescent males that navigate through vegetation at low light levels to locate their bioluminescent females, have significantly larger eyes than the males of diurnal species [35]. In contrast, diurnal males have significantly longer antennae than their conspecific females and usually also significantly longer antennae than the males of bioluminescent species [35], reflecting the importance of male antennae for the detection of pheromones in diurnal species.

It is currently unknown whether this antennal size dimorphism is also reflected in antennal sensilla diversity, and/or in the number and density of different sensilla types, including olfactory sensilla that are important for pheromone detection. We also presently do not know to what extent bioluminescent species retain the use of long-range pheromones used during mate search, (e.g., to lead males close enough to females, so both can detect and respond to the light signals of their potential mates). During firefly evolutionary history, there were several independent reversals from nocturnal activity with use of light signals to diurnal activity with pheromones as the main mating signal [36–38], and such reversals also took place several times within the genus *Photinus* [34]. This suggests that bioluminescent species likely retain the ability to use pheromones, at least to some degree, facilitating the reversal to exclusive pheromone use. Combinations of pheromone and bioluminescent signaling has been indicated at least once in the following genera; *Cyphonocerus*, *Pyrocoelia*, *Erthrolychnia*, *Phaenolis*, *Phausis*, *Pleotomus,* and *Lamprohiza* [33, 36, 37, 39, 40]. Field observations suggest that the nocturnal Blue Ghost firefly, *Phausis reticulata,* may use both light signals and pheromones during mate search [31], and the use of both pheromones and bioluminescence has been recently confirmed with field experiments for *Lamprohiza splendidula* [40]. In addition, the unusual diurnal, but bioluminescent, firefly species *Phosphaenus hemipterus* uses pheromones as the primary sexual signal and its faint bioluminescent glow as an aposematic defense signal, rather than for mating [41, 42].

While the bioluminescent mating signals of nocturnal species have been studied extensively, so far little is known about the pheromone signaling system in fireflies. The specific chemicals produced by female fireflies have only been identified for a hand full of species; *Pyrocoelia oshimana* [43], two unidentified *Diaphanes* species, *Pyrocoelia praetexta*, and *Lamprigera tenebrosa* [44]. However, functional female sex pheromones have only been identified for one species *Photinus corruscus* [21]. The specific antennal sensilla that respond to sex pheromones in *P. corruscus*, the diurnal winter firefly, have recently been identified as sensilla basiconica [21]. Once in close physical contact, both bioluminescent and non-bioluminescent fireflies will “antennate” each other intensively before mating, suggesting the sampling of contact chemicals, possibly cuticular hydrocarbons (CHCs) with gustatory receptors, for a final verification of a conspecific mate. South et al. [45] found CHCs on the pronotum and elytra of the diurnal species *P. corruscus* and *Lucidota atra,* but found only low or undetectable levels of CHCs in the nocturnal species *Photinus greeni*, *P. ignitus*, and *P. obscurellus* (it is unclear to what extent this may have been influenced by the different extraction methods used). *Photinus corruscus* males were able to distinguish between the CHCs of conspecific and heterospecific (*L. atra*) females, suggesting that CHCs are possibly used for reproductive isolation in this species [45].

In this study we document the sensilla diversity of both males and females of seven species of fireflies, including three diurnal and four nocturnal species. An interesting challenge was the identification of pheromone sensilla, especially in the absence of functional verification with electrophysiology. The first pheromone sensilla for fireflies were just recently identified in *P. corruscus* [21] and we used this opportunity to test our set of predictions for identifying potential sex pheromone sensilla in fireflies based on morphology alone. We hypothesized that the differences in mating signals between diurnal and nocturnal fireflies will be reflected in their antennal sensilla counts and possibly in their sensilla densities. Given the importance of pheromones for diurnal firefly species, we predicted to find (1) more chemoreceptors (including pheromone receptors) in diurnal species. If nocturnal species have completely lost the ability to use pheromones during mate search, we would expect to find (a) at least one sensilla type among the chemoreceptors that is only present in diurnal fireflies; if nocturnal species retain the ability to use pheromones we would expect (b) at least one sensilla type among the chemoreceptors that is present in greater numbers and possibly greater density on the antennae of diurnal species, compared to nocturnal species. We further predict that (2) males have more chemosensilla than females, and especially the diurnal males, that rely on tracking a pheromone plume to conspecific females, will have greater numbers, and possibly a greater density, of a pheromone chemoreceptor than their females. Finally, (3) pore or openings in the sensilla surface are necessary to allow volatile chemicals to enter the internal sensilla space and bind to receptor proteins on the sensory neurons [46], therefore any candidate for a pheromone sensilla should have pores. In the case of gustatory chemoreceptors that pick-up contact chemicals during antennation, we would not predict a difference between diurnal and nocturnal fireflies, since both diurnal and nocturnal species engage in antennation behavior before mating. We also would not predict a sex difference in gustatory chemoreceptors since both sexes likely use CHCs for mate recognition; this is suggested via the lack of sex differences in diurnal firefly CHC profiles [45]. In order to detect these contact chemicals, gustatory sensilla also should have pores. Based on the antennation behavior, gustatory chemoreceptors will possibly be concentrated on the ventral surface of the more distal antennomeres. Our proposed pheromone and gustatory chemoreceptor sensilla candidates will facilitate future functional testing of different sensilla morphotypes, an important next step towards understanding how fireflies use their antennae to perceive their world.

## Materials and Methods

### Taxon sampling

The seven firefly species in this study were selected to represent phylogenetic diversity (i.e., six genera in four subfamilies: Luciolinae, Lampyrohizinae, Photurinae, and Lampyrinae), sexual signal diversity (i.e., bioluminescent: glows or flashes and non-bioluminescent), and differences in dial activity (i.e., nocturnal and diurnal). The seven taxa in this study include four bioluminescent species (in four genera): three flashing species: *Photinus pyralis*, *Photuris lucicrescens,* Luciolinae sp. (an unidentified species from Africa in the Luciolinae subfamily), and one glowing species: *Phausis* sp. (a new species, discovered by Sarah Lower, that remains undescribed; voucher specimens KSH#8663 and KSH#8667 are stored in the Stanger-Hall lab at UGA), as well as three non-bioluminescent species (in three genera): *Lucidota punctata*, *Pyropyga nigricans*, and *Photinus corruscus* (formerly *Ellychnia corrusca* [48]). Species were determined to be bioluminescent or non-bioluminescent as adults by the presence or absence of a light organ on abdominal ventrites 5/6 or 6/7; flashing and glowing bioluminescent species were differentiated based on field observations and records in the literature (Table 1). The antennae of three males and three females of each species were examined. Vouchers are kept in the Stanger-Hall lab at the University of Georgia (Table S1).

**Table 1.**
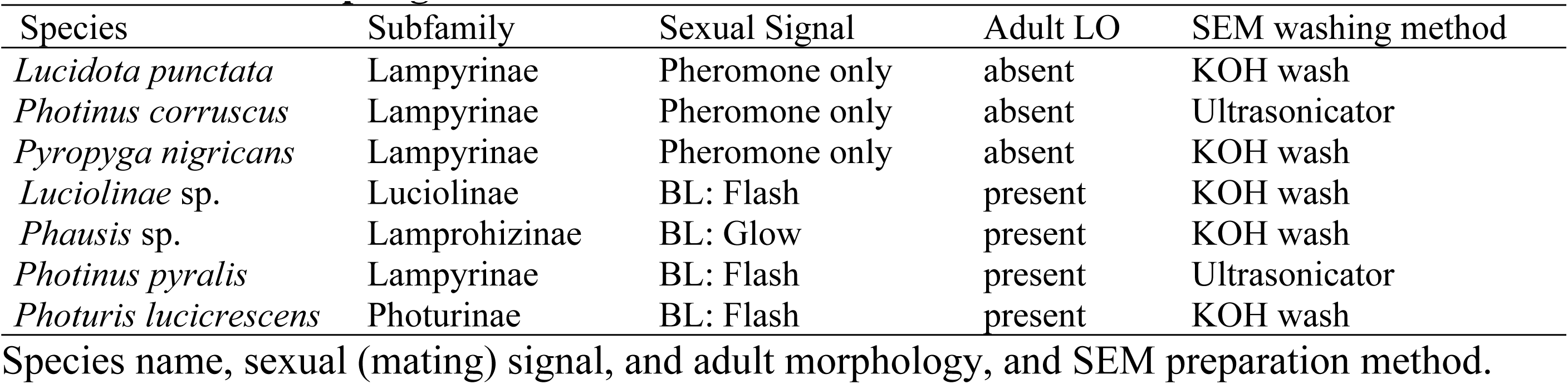
Taxon sampling.

### Scanning electron microscopy (SEM)

Specimens were prepared by removing both the left and right antennae from the head of each specimen. Antennae of the first several specimens were washed using soapy water and an ultrasonicator to remove debris. This method was effective for several specimens, but damaged others. Damaged specimens were not further examined and replaced with a new specimen, which was washed with a new method: antennae were placed in a 0.01% solution of KOH at 50° C for 2 hours. There was no difference in the structure of the antennal cuticle observed between the two cleaning methods. The antennae were then rinsed in distilled water and allowed to air dry for ∼10 minutes (both methods), before mounting them on SEM stubs, with one antenna facing dorsal side up and the other facing ventral side up, providing both a dorsal and a ventral view of the antennae for each individual firefly. Antennae were further air dried on their stubs for a minimum of 72 hours to ensure even sputter-coating. The antennae were sputter-coated with 30 nm of gold using the Lecia ACE600. Sputter-coated antennae were then imaged using the FE-SEM Thermo Fisher Teneo with an EDT and T1 detectors at the Georgia Electron Microscopy laboratory. For each individual antenna, SEM images were obtained for each individual antennomere. Each image contained a scale bar (generated by the Teneo software) for subsequent antennal area measurements.

### Antennal area

All taxa sampled in this study have filiform or moderately serrate (*Lucidota punctata*) antennae with 11 antennomeres (Fig 1), except *Phausis* sp. females which are paedomorphic (lacking both elytra and metathoracic wings but with paired pretarsal claws and a pair of stemmata [49] and have only three antennomeres. The area of each antennomere was measured (in mm^2^) using the polygon selection tool in ImageJ v1.52 [50] and the scale bars produced during imaging. The area of all (11 or 3) antennomeres was then summed for each specimen to determine the total antennal area for each side of the antenna. The area measurements for the dorsal and ventral sides of each individual antenna were then added to determine the total antennal area (per antenna) of each specimen. Since larger fireflies tend to have larger antennae, we examined the influence of body size on antenna size between our study taxa by measuring pronotum length as a proxy for body size [35] and used scaled antennal area (antenna area/pronotum length) to compare antennal areas between fireflies (note: due to the lack of pronotal expansions in Luciolinae, pronotum length will underestimate body size for Luciolinae compared to other fireflies). Pronotum length was measured from the anterior to the posterior edge along the midline of the pronotum. To visualize the relationship between pronotum length (body size) and total antenna area across the 4 nocturnal and 3 diurnal species in our study, we plotted antennal area against pronotum length for all 42 individuals in our analysis and labeled our samples as diurnal/nocturnal and male/female.

### Sensilla morphotypes, diversity and evolution

To describe sensilla morphotypes within and across our focal firefly species, we used key morphological characteristics as defined by Schneider [6] for different insect sensilla morphotypes. These characteristics included: shape of sensilla base (raised or not raised), shape of stalk (equal width throughout length or unequal width), length of stalk relative to base (stalk length equal to base or stalk length > 1x base length), and the presence or absence of grooves and/or pores. These morphological characteristics were used to separate sensilla morphotypes into the 9 major groups defined by Schneider [6]. In addition, we consulted Faucheux et al. [15] who examined the sensilla morphology of *Agriotes* (Elateridae) and synonymized sensilla nomenclature across Elateridae, a beetle family closely related to Lampyridae [51, 52]. We used the Elateridae naming scheme to classify each of the sensilla morphotypes for Lampyridae. Sensilla morphotypes that were not previously described or named, were named using the next consecutive number(s) of the Faucheux naming scheme [15]. We report all sensilla morphotypes found in males and females of each species. To identify a potential phylogenetic influence on the distribution of sensilla morphotypes in our seven study species and *Luciola cruciata* [23], we plotted all sensilla morphotypes on a cladogram of all 8 species (pruning all other species from the phylogeny of Martin et al. [51]).

### Sensilla diversity measures

Sensilla richness, or the number of sensilla types present, was counted for each specimen. To compare sensilla diversity across specimens and across firefly species we used the proportions of each sensilla type present (p_i_) on the antenna of a given specimen to calculate two different diversity indices that provide different measures of diversity. The general Simpson dominance index (D) is calculated as D = ∑(p_i_)^2^ and favors dominant sensilla types over rarer sensilla types. In contrast, the Shannon Index (H) is calculated as H=∑[-log(p_i_)] (p_i_) and favors rare sensilla types, allowing for a more even contribution of each sensilla type to the diversity index [53]. Since the Shannon and Simpson indices for different species cannot be directly compared or combined for analysis, we transformed all index values into their effective numbers (E) using E_D_ (Shannon)=10^H^ and E_H_ (Simpson)=1/D [53]. These effective numbers linearize values calculated from non-linear indices and represent how much more or less diverse in sensilla types one antenna is compared to another. We then used parametric t-Tests for E_D_ (normally distributed) and non-parametric Wilcoxon rank-sum tests for E_H_ (not normally distributed) to compare the average effective numbers (E_D)_ and E_H_) of each species to determine if there was a difference in sensilla diversity between diurnal and nocturnal species.

### Sensilla counts and sensilla density

For each specimen (three females and three males for seven species = 42 specimens) we counted the total number of sensilla for each sensilla morphotype on each antennomere (both dorsal and ventral side). To calculate the total sensilla number for each morphotype on one antenna we added the counts across the 11 antennomeres (three for *Phausis* females) of each individual specimen. We generated sensilla counts for two main functional categories (all mechanoreceptors and all chemoreceptors), as well as the total sensilla count per antenna. To determine how closely sensilla were packed on the antennal surface, we calculated sensilla density by dividing the total sensilla counts (per morphotype and category) by the total antennal area of the respective specimen. For summary statistics we used JMP Pro v17.2 [54] to calculate the mean and standard deviation (x̵±stdev) of sensilla numbers and sensilla densities for each sex and each species (all sensilla, all mechanoreceptor sensilla, and all chemoreceptor sensilla (Table 2, 3), and of individual sensilla morphotypes (Table S2, S3, S4, S5, S6). To examine how fireflies may optimize the sensitivity of their antennae for different stimuli, we plotted how total sensilla numbers and sensilla densities change on firefly antennae as antenna size increases.

**Table 2.** Mean sensilla counts and antennal areas of male and female fireflies.

| Species | Sex | Active | All sensilla (N) | All mechano-sensilla (N) | All chemo-sensilla (N) | Pronotum length (mm) | Antennal area (mm<sup>2</sup>) |
| --- | --- | --- | --- | --- | --- | --- | --- |
| *L. punctata* | F | D | 2535 ± 177 | 2133 ± 158 | 402 ± 21 | 1.3 ± 0.04 | 0.9 ± 0.1 |
| | M | D | 6984 ± 301 | 3405 ± 243 | 3579 ± 277 | 1.4 ± 0.2 | 2.3 ± 0.3 |
| *P. corruscus* | F | D | 5239 ± 381 | 4481 ± 559 | 757 ± 193 | 2.8 ± 0.1 | 1.6 ± 0.2 |
| | M | D | 5361 ± 572 | 4531 ± 454 | 830 ± 133 | 2.5 ± 0.2 | 1.6 ± 0.5 |
| *Py. nigricans* | F | D | 4487 ± 566 | 3665 ± 388 | 822 ± 279 | 1.4 ± 0.3 | 1.1 ± 0.4 |
| | M | D | 3978 ± 1141 | 2800 ± 671 | 1178 ± 470 | 1.5 ± 0.3 | 1.5 ± 0.9 |
| Luciolinae sp. | F | N | 2143 ± 152 | 1675 ± 135 | 468 ± 16 | 1.4 ± 0.1 | 0.5 ± 0.05 |
| | M | N | 2307 ± 131 | 1789 ± 68 | 517 ± 29 | 1.1 ± 0.1 | 0.6 ± 0.02 |
| *Phausis* sp. | F | N | 49 ± 7 | 39 ± 4 | 10 ± 4 | 0.8 ± 0.06 | 0.02 ± 0.00 |
| | M | N | 1131 ± 114 | 559 ± 115 | 572 ± 24 | 0.9 ± 0.1 | 0.2 ± 0.04 |
| *P. pyralis* | F | N | 3708 ± 285 | 2997 ± 190 | 711 ± 99 | 1.9 ± 0.1 | 2.0 ± 0.6 |
| | M | N | 3937 ± 354 | 3034 ± 292 | 903 ± 151 | 1.8 ± 0.3 | 2.5 ± 0.2 |
| *Ph. lucicrescens* | F | N | 2787 ± 164 | 1947 ± 116 | 840 ± 54 | 3.2 ± 0.2 | 2.6 ± 0.6 |
| | M | N | 2710 ± 444 | 1988 ± 118 | 722 ± 240 | 2.8 ± 0.06 | 1.9 ± 0.2 |

Sensilla counts (mean ± stdev) per antenna for all sensilla types, all mechanoreceptors, all chemoreceptors, mean pronotum length, and mean antennal area, by species and sex (F: 3 females, M: 3 males, D: diurnal, N: Nocturnal, *L.* = *Lucidota*, *P.* = *Photinus*, *Py.* = *Pyropyga*, *Ph.* = *Photuris*).

### Sensilla distribution

To assess whether different parts of the antennae could be used for different functions (e.g., mechanoreception, chemoreception), we determined how the different sensilla morphotypes were distributed across the length of the antenna of each individual specimen. To visualize this distribution of the sensilla in each functional group and each morphotype, we plotted the mean number of sensilla per antennomere for each sex and each species across the 11 antennomeres.

### Statistical analyses

We used a two-way mixed model ANOVA in JMP Pro v17.2 to test for differences in antenna size, sensilla counts, and sensilla densities between diurnal and nocturnal fireflies and between males and females. Due to our small sample sizes within firefly species (3 males and 3 females each) we used “species” as a random effect (covariate) for all our analyses, with activity and sex as fixed effects. Since males and females of diurnal and nocturnal taxa were predicted to differ for some sensilla types, we also tested for an interaction between sex and activity. For each mixed model analysis, we report the amount of variation explained by the species covariate. All fixed effects tests were based on 1 parameter (Nparm=1) and 1 degree of freedom for the numerator (DFNum), therefore we only report the degrees of freedom for the denominator (DFDen = error term), the F-ratio (F) for testing that the effect is zero, and the p-value (Prob>F) for the fixed effects (and their interaction) for each analysis. When the interaction term was significant, we tested effect slices (activity by sex, sex by activity) with Student’s t tests to test our predicted differences in chemoreceptors (but not mechanoreceptors) between diurnal and nocturnal males, and between diurnal males and diurnal females. Since p-values represent a continuum, we report p-values with p<0.05 as statistically significant, and p-values close to 0.05 as marginally significant.

Since larger fireflies tend to have larger antennae [35], we used a separate mixed model analysis to examine the distribution of body sizes (pronotum length) across our firefly sample and report the percent variation explained by species (random effect). To specifically examine the influence of body size on antenna size across our specimens, we added pronotum length as an additional (continuous) fixed effect to a mixed model analysis with activity and sex (and sex*activity) as fixed effects and species as a random effect.

## Results

### Antenna size

The antennal areas across our seven species of fireflies varied greatly (Table 2). The smallest antennal area (x̵±stdev) was measured in *Phausis* sp. females (three antennomeres: x̵_Female_=0.02mm^2^ ± 0.001) and males (11 antennomeres: x̵_Male_ = 0.2mm^2^ ± 0.04). The females of nocturnal *Photuris lucicrescens* (x̵_Female_ = 2.6mm^2^ ± 0.6) and *Photinus pyralis* (x̵_Female_ = 2.5mm^2^ ± 0.2) had the largest measured antennal areas (Table 2), however, when body size was taken into account (scaled antenna size=antennal area/pronotum length), the males of diurnal *Lucidota punctata* and nocturnal *P. pyralis* had the largest (scaled) antennal areas (Table 3).

**Table 3:** Mean sensilla densities and scaled antennal areas of male and female fireflies.

| Species | Sex | Activity | All sensilla<br>(N/mm<sup>2</sup>) | All mechano-<br>sensilla<br>(N/mm<sup>2</sup>) | All chemo-<br>sensilla<br>(N/mm<sup>2</sup>) | Scaled antennal<br>area<br>(area/pronotum<br>length mm) |
| --- | --- | --- | --- | --- | --- | --- |
| *L. punctata* | F | D | 2889 ± 454 | 2431 ± 394 | 458 ± 59 | 0.7 ± 0.08 |
|  | M | D | 3020 ± 524 | 1467 ± 208 | 1553 ± 334 | 1.7 ± 0.4 |
| *P. corruscus* | F | D | 3294 ± 159 | 2810 ± 196 | 483 ± 155 | 0.6 ± 0.05 |
|  | M | D | 3416 ± 577 | 2892 ± 516 | 524 ± 67 | 0.6 ± 0.2 |
| *Py. nigricans* | F | D | 4277 ± 998 | 3420 ± 604 | 857 ± 473 | 0.8 ± 0.1 |
|  | M | D | 3438 ± 1834 | 2435 ± 1319 | 1002 ± 536 | 0.9 ± 0.4 |
| Luciolinae sp. | F | N | 3927 ± 111 | 2431 ± 394 | 458 ± 59 | 0.4 ± 0.007 |
|  | M | N | 5023 ± 41 | 3896 ± 76 | 1127 ± 55 | 0.4 ± 0.02 |
| *Phausis* sp. | F | N | 2671 ± 256 | 2127 ± 71 | 544 ± 207 | 0.02 ± 0.002 |
|  | M | N | 5955 ± 803 | 2897 ± 174 | 3058 ± 697 | 0.2 ± 0.002 |
| *P. pyralis* | F | N | 1887 ± 408 | 1526 ± 331 | 360 ± 83 | 1.05 ± 0.3 |
|  | M | N | 1604 ± 50 | 1235 ± 16 | 369 ± 61 | 1.4 ± 0.1 |
| *Ph. lucicrescens* | F | N | 1090 ± 235 | 761 ± 158 | 330 ± 76 | 0.8 ± 0.1 |
|  | M | N | 1399 ± 101 | 1031 ± 34 | 369 ± 87 | 0.7 ± 0.8 |

Sensilla density (mean ± stdev) per antenna for all sensilla types, all mechanoreceptors, all chemoreceptors, mean antennal area scaled by pronotum length by species and sex. (F: 3 females, M: 3 males, D: diurnal, N: Nocturnal, *L.* = *Lucidota*, *P.* = *Photinus*, *Py.* = *Pyropyga*, *Ph.* = *Photuris*).

### Body size and antennal area

Our species sample varied greatly in body size (pronotum length; Table 2). In our mixed model analysis, species (covariate) accounted for 95.06% of the variation in body size. Body sizes in our species sample did not differ significantly by activity or sex (or sex*activity). Overall, the antennal areas of the 42 firefly specimens in our study were positively correlated (R= 0.467; Spearman’s rho =0.668, p<0.0001) with body size (Fig 2A), which means larger fireflies tend to have larger antennae.

**Fig 2.**
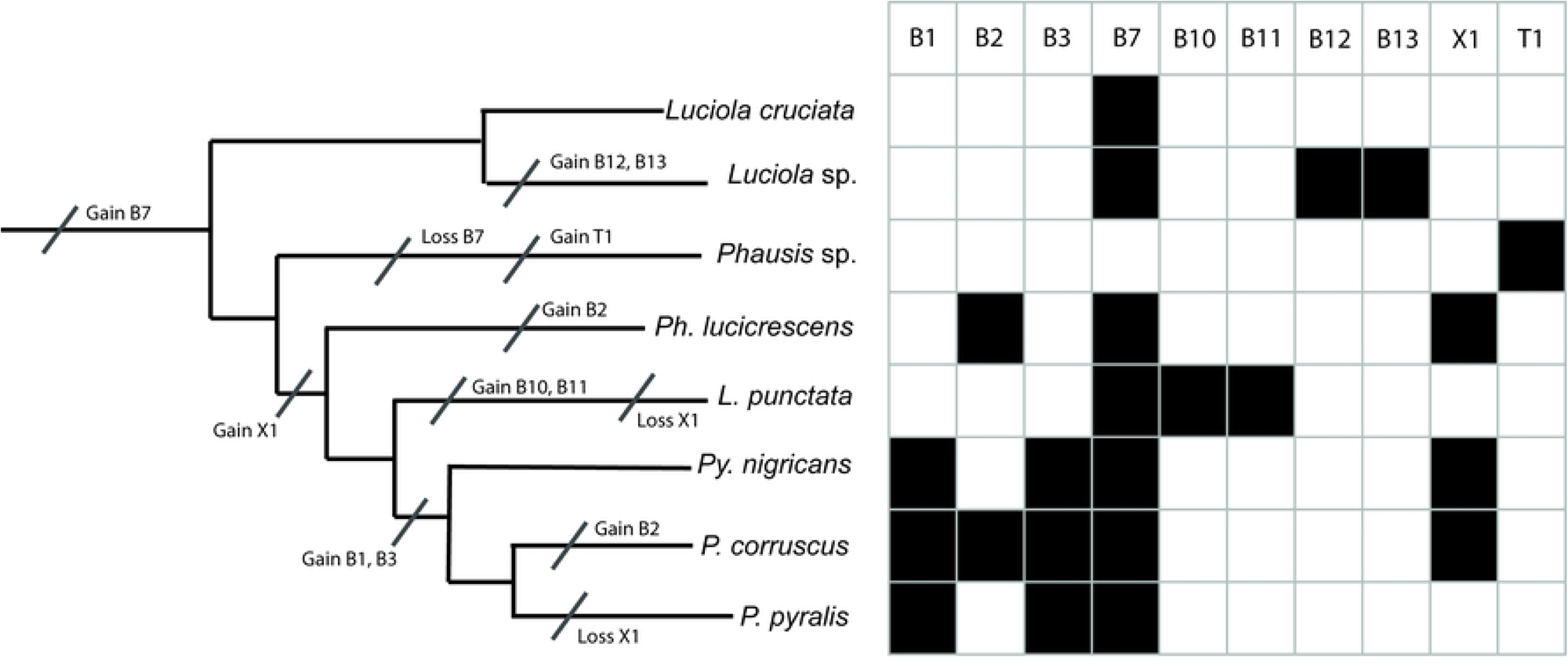
Firefly antennal area. (A) Relationship between body size (pronotum length) and antennal area: antennal area is positively correlated with body size (R= 0.467; Spearman’s rho =0.668, p<0.0001); (B) Comparison of antennal area by activity time: there is no difference in antennal size between diurnal and nocturnal species (x̵_Diurnal_ =1.50±0.6 mm^2^, x̵_Nocturnal_ =1.28±1.07 mm^2^, DFDen= 5.0, F=0.077, p=0.7943); (C) Comparison of antennal area by sex: males have significantly larger antennae than females (x̵_Male_ =1.49±0.9 mm^2^, x̵_Female_ =1.26± 0.9 mm^2^; DFDen=33.4, F=11.58, p=0.0017).

In our mixed model analysis of antenna size, species accounted for 77.34% of the variation in antennal area. Body size (pronotum length) had a significant effect on antennal area (DFDen=17.0, F=16.65, p=0.0008). The antennal areas of the diurnal and nocturnal (x̵_Diurnal_ =1.50±0.6 mm^2^, x̵_Nocturnal_ =1.28±1.07 mm^2^) fireflies in our study did not differ significantly from each other (mixed model, DFDen= 5.0, F=0.077, p=0.7943, Fig 2B), but males had significantly larger antennal areas than females (x̵_Male_=1.49±0.9 mm^2^, x̵_Female_ =1.26± 0.9 mm^2^; DFDen=33.4, F=11.58, p=0.0017, Fig 2C). The interaction term (activity*sex) was also significant (DFDen=32.2, F=5.39, p=0.0267) with significantly larger antennal areas in diurnal males compared to diurnal females (x̵_Male_=1.812± 0.66 mm^2^, x̵_Female_=1.2±0.38 mm^2^; Student’s t ratio = - 3.86, p=0.0005, Fig S1).

### Sensilla morphology

We identified a total of 14 different sensilla morphotypes across the seven firefly species. Two of these 14 morphotypes were first identified in *L. cruciata* [23]. One morphotype was extremely rare and was not counted, two other morphotypes were found in only one sex, or in only a few specimens within a species. Eleven morphotypes, if present at all, were present in both sexes (Table 4). Based on their morphology [6, 15], we identified 3 mechanoreceptor and 8 chemoreceptor morphotypes among the 11 most common sensilla morphotypes in our 7 species (based on morphology, the three rare morphotypes are likely chemoreceptors).

**Table 4.** Distribution of sensilla morphotypes across species.

|  | *P. corruscus* | *Py. nigricans* | *L. punctata* | *P. pyralis* | *Ph. lucicrescens* | *Phausis* sp. | Luciolinae<br>sp. | *L. cruciata*<sup>1</sup> |
| --- | --- | --- | --- | --- | --- | --- | --- | --- |
| Sensilla<br>type | Diurnal | Diurnal | Diurnal | Nocturnal | Nocturnal | Nocturnal | Nocturnal | Nocturnal |
| C1 | + | + | + | + | + | + | + | + |
| C2 | + | + | + | + | + | + | + | - |
| SC | + | + | + | + | + | - | + | - |
| B1 | + | + | - | + | - | - | - | - |
| B2 | + | - | - | - | + | - | - | - |
| B3 | + | + | - | + | - | - | - | - |
| B7 | + | + | + | + | + | - | + | + |
| B10 | - | - | + | - | - | - | - | - |
| B11 | - | - | + | - | - | - | - | - |
| B12 | - | - | - | - | - | - | +<sup>2</sup> | - |
| B13 | - | - | - | - | - | - | + | - |
| X1 | + | + | - | - | + | - | - | - |
| T1 | - | - | - | - | - | + | - | - |

Presence (+) or absence (-) of 13 sensilla morphotypes in eight species of Lampyridae. (*L.* = *Lucidota*, *P.* = *Photinus*, *Py.* = *Pyropyga*, *Ph.* = *Photuris*). <sup>1</sup>*L. cruciata* data from Iwasaki et al., 1995; <sup>2</sup>B12 sensilla are only present in male *Luciola* sp.)

#### Mechanoreceptors

<u>Sensilla chaetica</u>, also known as the bristle or spine sensilla (Schneider 1967), are slender sensilla arising directly from the antennal cuticle, with no modified ring around the base. These sensilla vary in length, but all gradually taper towards the distal end. The walls of these sensilla can be smooth or bear grooves that run parallel to the length of the stalk. We found two sensilla chaetica variants in fireflies (Fig 3). (1) <u>Sensilla chaetica type 1</u> (C1, Fig 3A): C1 are long bristle sensilla with grooves running parallel lengthwise along the stalk. These are the longest sensilla found on firefly antennae and were located on all 11 antennomeres. They were distributed evenly across the surface of each individual antennomere. C1 sensilla were found in all seven species examined. (2) <u>Sensilla chaetica type 2</u> (C2, Fig 3B): C2 sensilla have a short smooth stalk that comes to a distinct point. These sensilla were found in all seven species examined.

**Fig 3.**
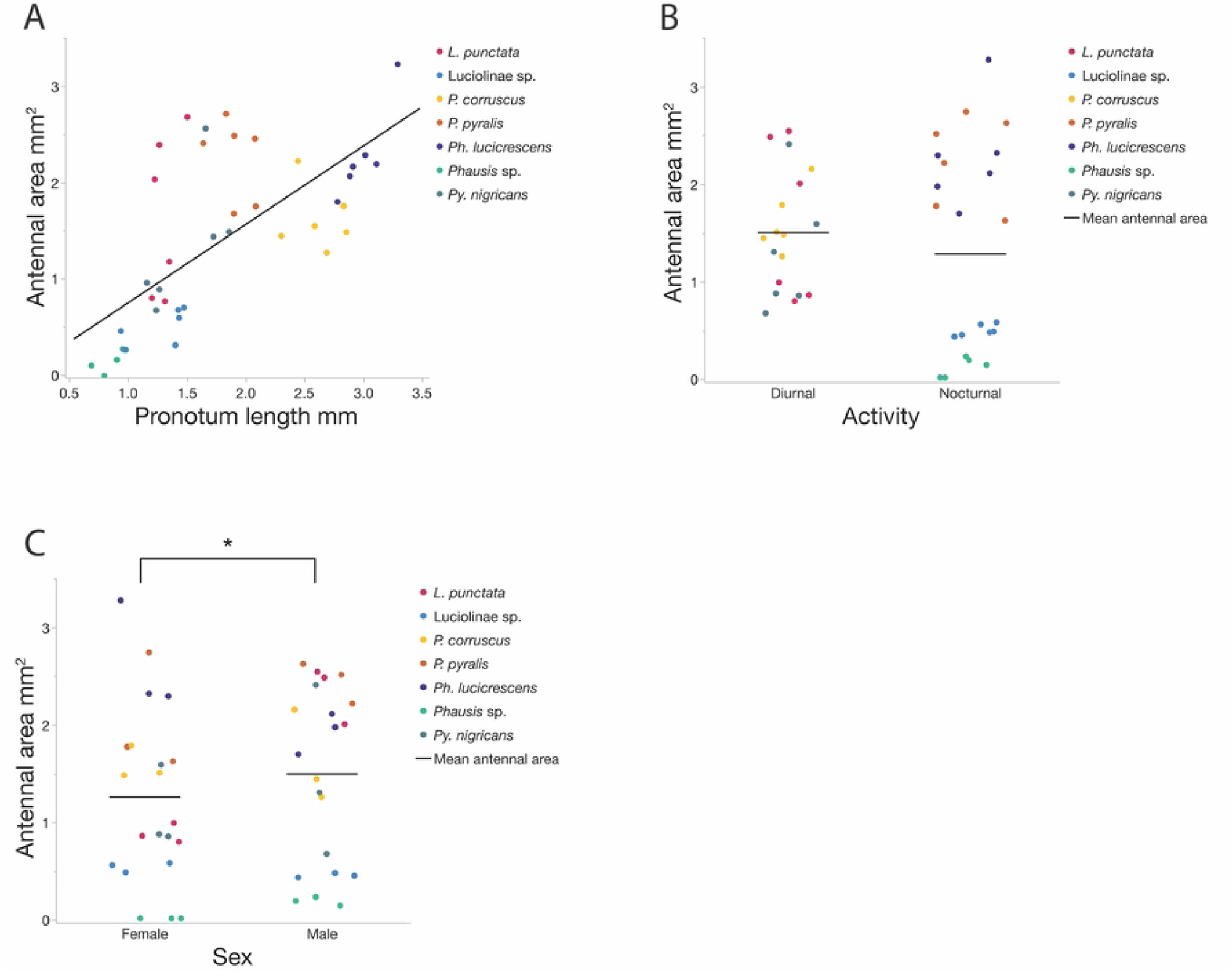
Mechanosensilla morphotypes. (A) mechanosensilla chaetica type 1 (C1) on the antenna of a *Py. nigricans* female (also pictured are cuticular pores (CP) and chemosensilla basiconica type B3); (B) mechanosensilla chaetica type 2 (C2) of a *Py. nigricans* male; (C) mechanosensilla campaniform sensilla (SC) of a Luciolinae sp. female (also pictured are chemosensilla basiconica type B13).

<u>Sensilla campaniform</u>. We identified one type of sensilla campaniform (SC; Fig 3C). These sensilla consist of a ring slightly raised above the antennal surface, with a raised disc, equal in height, inside the center of the ring. SC sensilla were found in low numbers relative to the other mechanoreceptors (Table 4). They were found on the antennae of all species except *Phausis* sp.; they were absent in male *Lucidota punctata* and female *Photuris lucicrescens* (Table 4).

#### Chemoreceptors

<u>Sensilla basiconica</u>, also known as peg or cone sensilla, are characterized by their raised base with a peg or cone arising from that base. The length and width of the peg varies between morphotypes. Peg and cone walls may or may not have pores and may have grooves that run parallel along the length of the peg (Schneider 1967). Basiconica sensilla were found in all seven firefly species and we identified six different variants (types): (1) <u>Sensilla basiconica type 1</u> (B1: Fig 4A) are peg sensilla with a well-defined dome shaped base and a short peg with a single pore at the distal end (Fig 5A). The peg length is ∼3-4 times the height of the dome base, the width of the peg is uniform throughout the length and the distal end of the peg is rounded. We found no pores along the length of B1 sensilla, however B1 sensilla identified by Lower et al. [21] possessed pores along the sides of the peg. B1 sensilla were found in the diurnal (non-bioluminescent) species *P. corruscus*, *Py. nigricans*, and the nocturnal (bioluminescent) species *P. pyralis*. (2) <u>Sensilla basiconica type 2</u> (B2: Fig 4A) are peg sensilla with a well-defined dome base and a moderate length peg (∼4-5 times the height of the dome base). The peg is widest at the base and gradually tapers in width coming to a blunt point at the distal end, no visible pores were found on the peg. B2 sensilla were found in the diurnal species *P. corruscus*, and the nocturnal species *Ph. lucicrescens* and Luciolinae sp. (3) <u>Sensilla basiconica type 3</u> (B3: Figs 3A, 4B, 5E) are peg sensilla with a modified dome base, with a flattened top, and a peg (∼5 times the height of the base). The width of the peg is equal throughout its length and bears pores in high density along the distal 3⁄4 (Fig 5B). The peg comes to a blunt point at its distal end. B3 sensilla were found in the diurnal species *P. corruscus*, *Py. nigricans,* and the nocturnal species *P. pyralis*. (4) <u>Sensilla basiconica type 7</u> (B7: Fig 4C) are peg sensilla with a dome base and peg (∼2-3 times the height of the base). The peg is equal in width for the first 2/3 of its length and then tapers to a distinct point. The last 1/3 of the peg has parallel grooves that run along the length of the peg and meet at the distal point. There were no visible pores on the peg of B7 sensilla. B7 sensilla were found in all studied species except *Phausis* sp. (5) <u>Sensilla basiconica</u> <u>type 10</u> (B10: Fig 4D, E) are cone sensilla with a raised collared base and a broad cone. The length of the cone is 2 times the height of the collared base, pores are present on the cone (Fig 5C). The cone is equal in width to the collared base for the first ½ of the length and then tapers gradually to a point. B10 sensilla were only found in the diurnal species *L. punctata*. (6) <u>Sensilla</u> <u>basiconica type 11</u> (B11: Fig 4E) are modified cone sensilla with a collared base and a rounded cone. The length of the cone is ∼1.5-2 times the length of the base. The proximal end of the cone is equal in width to the base, the sides then gradually diverge, increasing the width of the cone, with a rounded end. The distal end of the cone is broadly rounded to a shallow dome shape, the cone lacks visible pores. B11 sensilla were only found in the diurnal species *L. punctata*. (7) <u>Sensilla basiconica type 12</u> (B12: Fig 4F, G) have a membranous base with a short peg that gradually comes to a point at the distal end which bears a single pore. The length of the peg is about equal to the width at the base; these sensilla were only found in the males of Luciolinae sp. (8) <u>Sensilla basiconica type 13</u> (B13: Figs 3C and 4H) closely resemble B3 sensilla, but completely lack pores; these sensilla were only found in Luciolinae sp.

**Fig 4.**
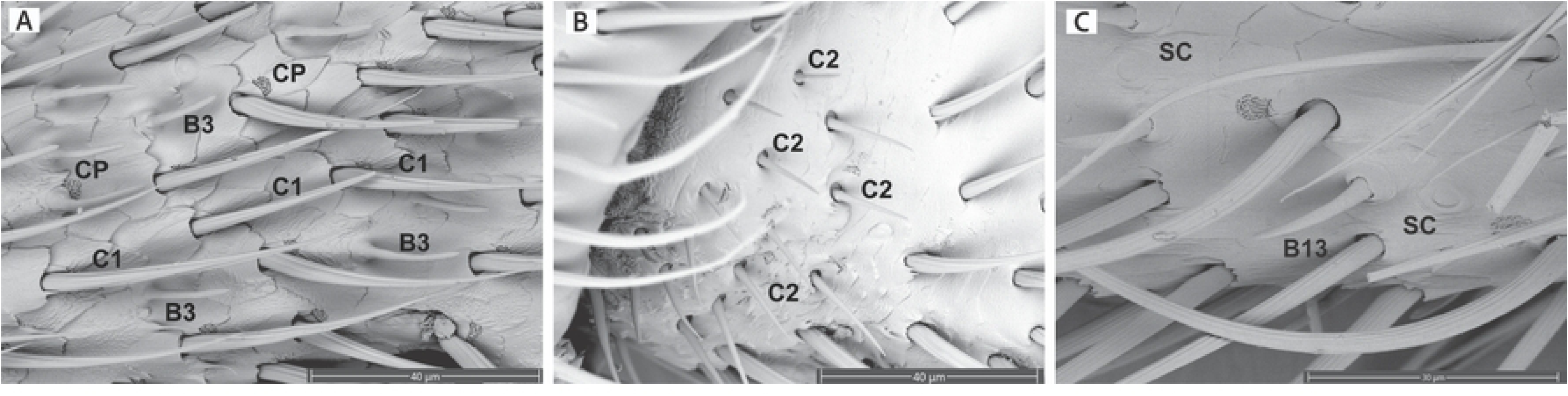
Chemosensilla. (A) sensilla basiconica types 1 and 2 (B1 and B2) on the antenna of a *P. corruscus* female; (B) sensilla basiconica type 3 (B3) of a *P. corruscus* male; (C) sensilla basiconica type 7 (B7) of a *P. corruscus* male; (D) sensilla basiconica type 10 (B10) of a *L. punctata* female; (E) sensilla basiconica types 10 and 11 (B10 and B11) of a *L. punctata* male; (F) sensilla basiconica type 12 (B12) and cuticular pores (CP) on a Luciolinae sp. female; (G) sensilla basiconica type 12 (B12) with an apical pore (arrow) of a Luciolinae sp. female; (H) basiconica sensilla types 7 and 13 (B7, B13) and cuticular pores (CP) of a Luciolinae sp. female; (I) sensilla trichodea (T1) of a *Phausis* sp. male.

**Fig 5.**
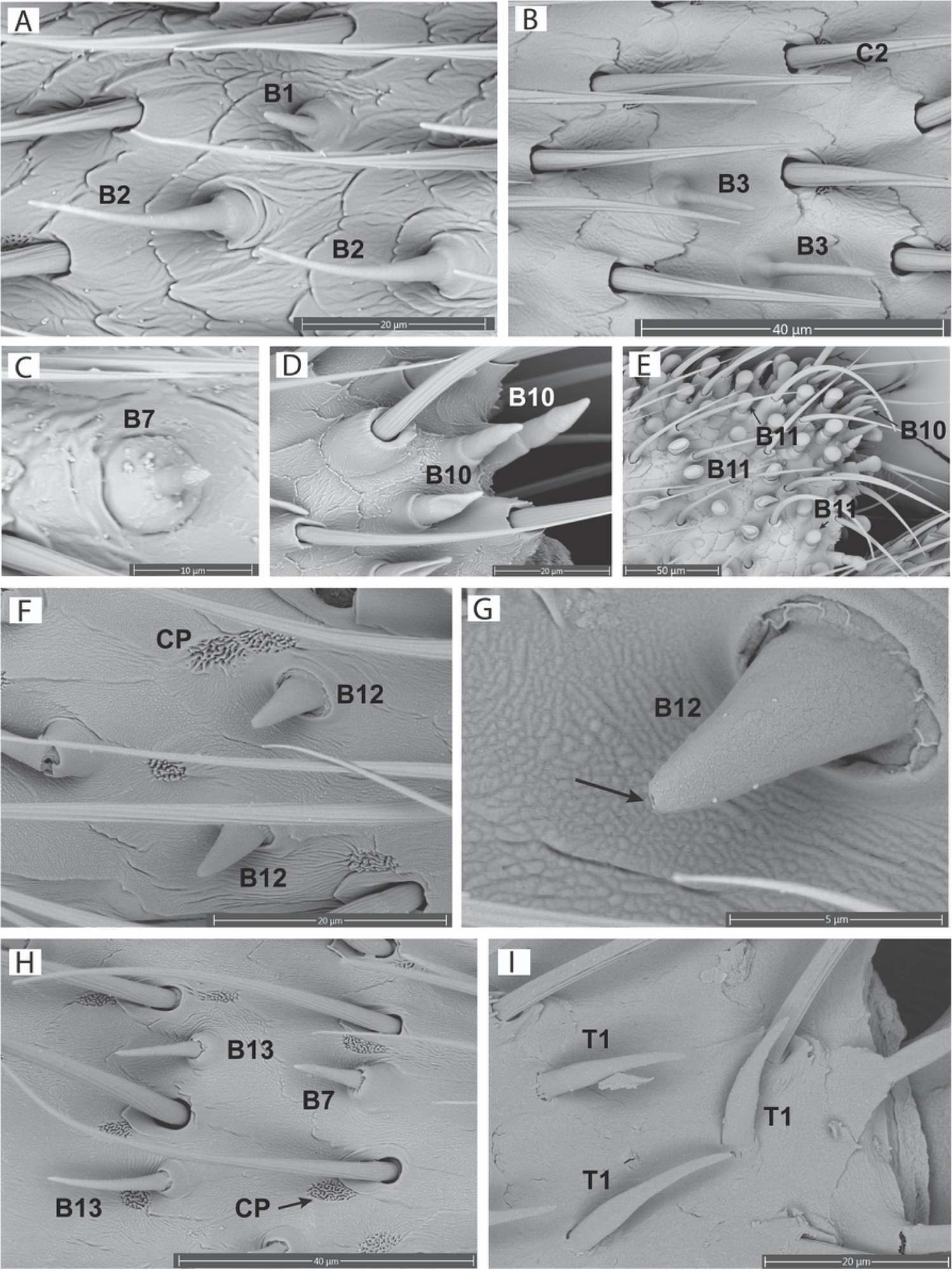
Sensilla pores and unique sensilla. (A) B1 sensilla with a single pore (arrow) and the sensilla peg slightly compressed into the dome base, on the antenna of a *P. corruscus* female; (B) multipored (arrows) peg of B3 sensilla of a *P. corruscus* male; (C) multipored (arrows) cone of B10 sensilla of a *L. punctata* male; (D) sensilla coeloconica (X1) of a *P. corruscus* female; (E) cone sensilla, surrounded by (broken) B3 sensilla on the antenna of a *P. corruscus* female.

<u>Sensilla coeloconica</u> (X1: Fig 5D) consist of pegs that arise from inside a pit, a rounded depression in the cuticle, therefore the proximal end of the peg is not visible inside the pit. The sensilla coeloconica in Lampyridae are characterized by a circular pit with a B7 type peg that comes to a point at the distal end with multiple longitudinal grooves meeting at the end, and no visible pores.

<u>Sensilla trichodea</u> (T1: Fig 4I) are simple hair like sensilla without a modified base, directly arising from the cuticle surface. We found only one type of sensilla trichodea (T1) in fireflies and exclusively in the nocturnal species *Phausis* sp. These sensilla lack pores and are short, with the width of the stalk at the base ∼8 times the width of the stalk at the distal end. The width of the sensilla gradually decreases before coming to a blunt point.

In addition, we found <u>cone sensilla</u> (Fig 5E) in very low numbers and not consistently present across all individuals of a species. These sensilla were seen in one or two individuals per species, with 1-3 cones present per individual. Cone sensilla have no raised base (differentiating them from basiconica sensilla) and are shaped like cones with the width at the base equal to ∼1.5-2 times the height; pores were not observed. They were found on *Photinus corruscus*, *Pyropyga nigricans*, and Luciolinae sp. antennae, but not in all specimens of any species. We also found clusters of large cuticular pores (CP: Figs 3A and 4F) on the antennal surface of the Luciolinae sp., *Photinus pyralis, Phausis* sp., and *Pyropyga nigricans*. These pores were not associated with sensilla but could often be found at the base of C1 sensilla.

### Sensilla diversity across species

We documented between three and eight sensilla morphotypes within an individual firefly species. The distribution of individual morphotypes varied greatly between species. Overall, three different mechanoreceptor sensilla types (sensilla chaetica C1, C2 and sensilla campaniform SC), nine different chemoreceptor sensilla types (sensilla basiconica types B1, B2, B3, B7, B10, B11, B12, B13, and sensilla trichodea T1), as well as one potential temperature/humidity receptor sensilla type (sensilla coeloconica X1), were identified across the seven firefly species in this study. Two mechanoreceptor types (sensilla chaetica types C1 and C2) were the only sensilla morphotypes found in all seven firefly species (Table 4). Among the chemoreceptors the B7 sensilla type was shared by most species (except *Phausis* sp.) and found in both males and females (Table 4). Other chemoreceptor sensilla were shared by three firefly species (B1, B3), by two species (B2) or they were unique for a single species (B10, B11: *Lucidota punctata*; B12, B13: Luciolinae sp.; T1: *Phausis* sp.). A potential thermo/hygroreceptor sensilla type (X1) was shared by three firefly species (Table 4). There was no single chemoreceptor morphotype found exclusively in diurnal species, or shared by all diurnal and nocturnal species, and the different chemoreceptor types varied greatly between species (Table 4). Our comparisons of sensilla diversity based on effective numbers for Shannon and Simpson diversity indices showed no significant difference in antennal sensilla diversity between diurnal and nocturnal fireflies, no matter whether rare (Shannon’s index effective numbers, t-Test; x̵_Diurnal_=2.37±0.53, x̵_Nocturnal_=2.37±0.86, df=13, p=0.99) or common (Simpson’s index effective numbers, Wilcoxon rank-sum, x̵_Diurnal_ =1.62±0.41, x̵_Nocturnal_ =1.95±0.47, df=13, p=0.061) sensilla types were favored.

### Sex differences in sensilla types

Almost all sensilla morphotypes were found in both males and females of the respective species, except for SC and B12 sensilla. SC sensilla were found in both sexes for the two *Photinus* species, *Pyropyga nigricans*, and Luciolinae sp., but they were only found in *Lucidota punctata* females and *Photuris lucicrescens* males. B12 sensilla were exclusively found in Luciolinae sp. males (Table 4).

### Evolution of sensilla morphotypes

Using the phylogeny of Martin et al. [51] to extract the evolutionary relationships of the eight firefly species in six genera studied to date, it is possible to hypothesize the ancestral pattern of sensilla. B7 appears to be the ancestral sensillum for the common ancestor of all 8 firefly species and was subsequently lost in *Phausis* (Fig 6). Both B1 and B3 sensilla most likely evolved in the common ancestor of *Pyropyga* and *Photinus* fireflies. Other sensilla types evolved independently in individual lineages, e.g., B2 evolved independently in *Photuris lucicrescens* and *Photinus corruscus*, B10 and B11 evolved within *Lucidota,* B12 and B13 evolved within Luciolinae sp., and T1 evolved within *Phausis*. X1 evolved independently in *Photuris*, *Pyropyga* and *P. corruscus;* alternatively, it could have evolved in *Photuris* and the common ancestor of *Pyropyga* and *Photinus* and was subsequently lost in *Photinus pyralis.* For *Luciola cruciata*, unique pored sensilla chaetica, capitular sensilla and gemmiform sensilla are reported in the literature [23].

**Fig 6.**
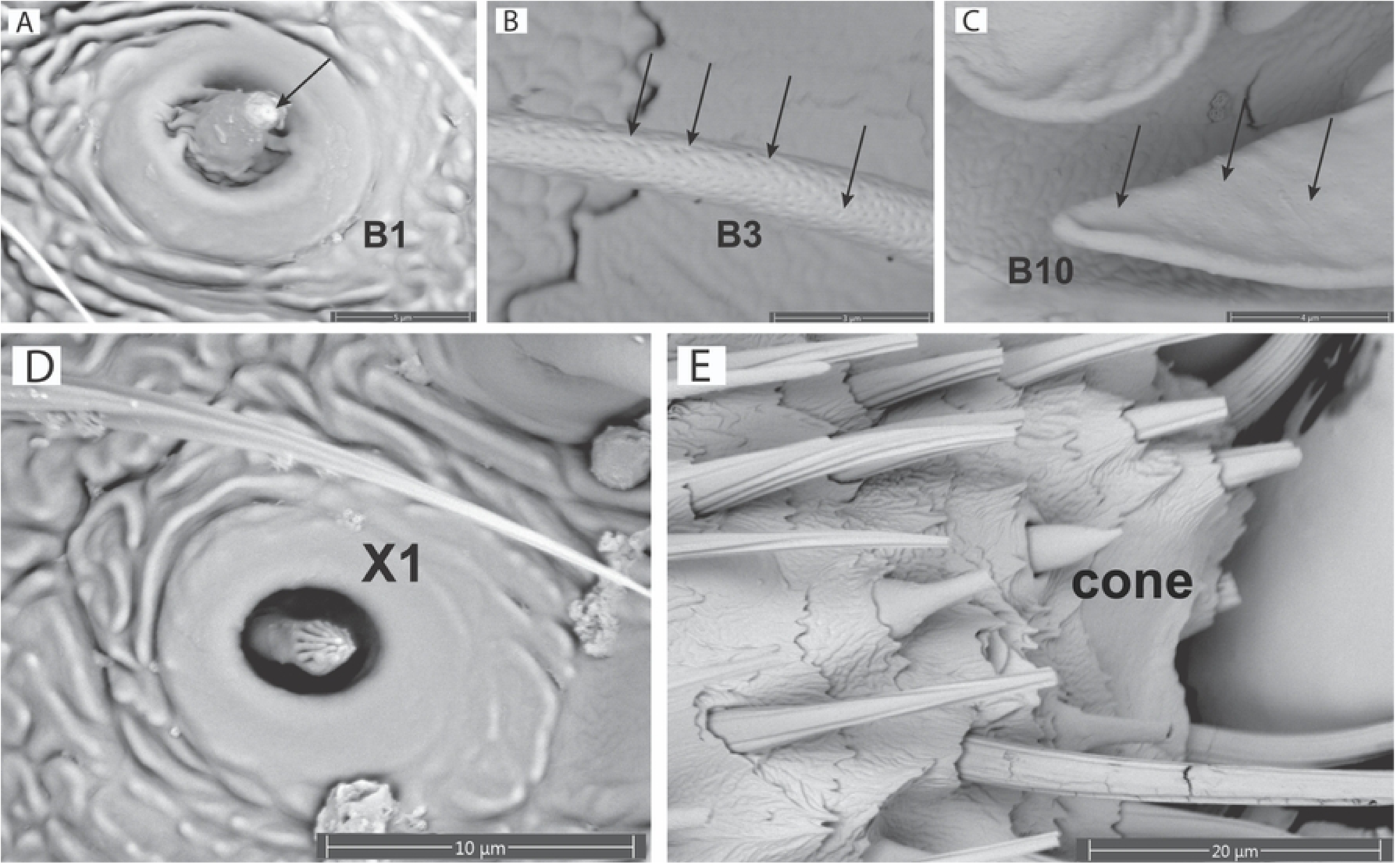
Evolution of sensilla morphotypes. Sensilla presence (black square) and absence (empty square) for each chemosensilla type across study species are marked in the data matrix. The most parsimonious gains and losses for each morphotype are marked with a hatchmark on the corresponding branches of the species cladogram.

### Sensilla counts and densities

The total antennal sensilla counts (x̵ ± stdev) for a single firefly antenna ranged from 49 ± 7 total sensilla per antenna in *Phausis* sp. females to 6984 ± 301 sensilla in *Lucidota punctata* males (Table 2). Sensilla densities in our species sample ranged from an average of 1090 ± 235 per mm^2^ in *Photuris lucicrescens* females to an average of 5955 ± 803 per mm^2^ in *Phausis* sp. males (Table 3). Total sensilla counts were positively correlated with antennal area across the 42 antennae of the seven firefly species (N=42, R^2^ =0.389, Spearman’s rho =0.6677, p<0.0001, Fig 7A), which means larger antennae had more sensilla. In contrast, sensilla densities were negatively correlated with antennal area (R^2^ = 0.542, Spearman’s rho= -0.7094, df = 1, p<0.0001, Fig 7B), which means that larger antennae had fewer sensilla per area than smaller antennae.

**Fig 7.**
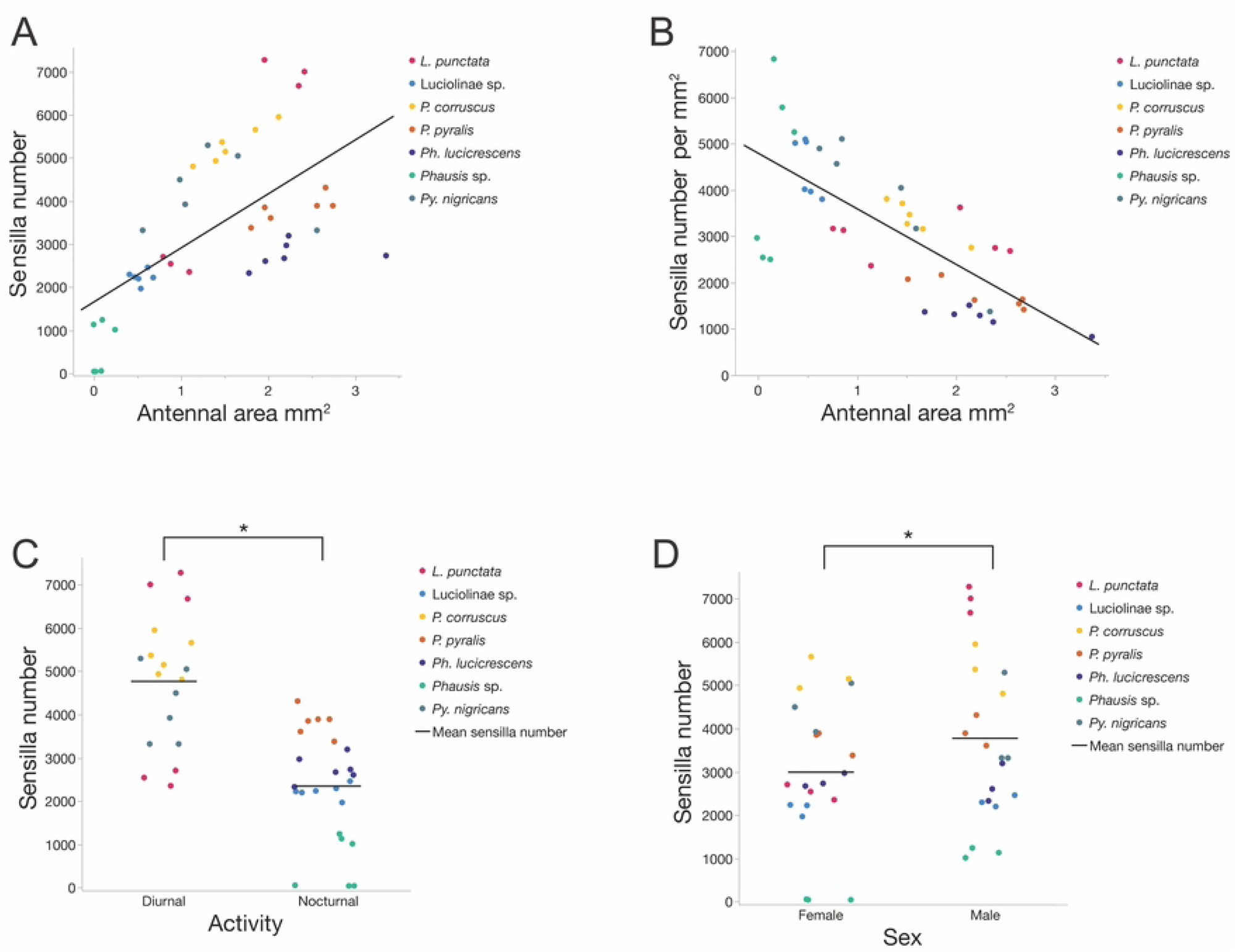
Antennal area and sensilla abundance by sex and activity. (A) Positive correlation between antennal area and total sensilla number (N=42, R^2^ =0.389, Spearman’s rho =0.6677, p<0.0001): larger antennae tend to have more sensilla; (B) Negative correlation between antennal area and total sensilla density (N-42, R^2^ = 0.542, Spearman’s rho= -0.7094, df = 1, p<0.0001): larger antennae tend to have a lower sensilla density; (C) Comparison of mean sensilla number between diurnal and nocturnal species: diurnal species have significantly more sensilla than nocturnal species (x̵_Diurnal_=4768±1495, x̵_Nocturnal_=2349±1242; DFDen=5, F=8.33, p=0.0343); (D) Comparison of mean sensilla number between females and males: males have significantly more sensilla than females (x̵_Male_=3776±1919, x̵_Female_=2995±1638; DFDen=38, F= 8.7, p=0.0054).

In our mixed model analysis, species (random effect covariate) accounted for 55.2% of the variation in total sensilla counts across firefly antennae. Diurnal fireflies had significantly more antennal sensilla than nocturnal fireflies (x̵_Diurnal_=4768±1495, x̵_Nocturnal_=2349±1242; DFDen=5, F=8.33, p=0.0343; Fig 7C) and males had significantly more sensilla than females (x̵_Male_=3776±1919, x̵_Female_=2995±1638; DFDen=38, F= 8.7, p=0.0054; Fig 7D). The interaction (activity*sex) was not significant. Similarly, species accounted for 69.5% of the variation in sensilla densities. Diurnal and nocturnal fireflies did not differ significantly (x̵_Diurnal_ ==3392±907 per mm^2^, x̵_Nocturnal_ =2949±1766 per mm^2^; DFDen=5, F=0.187, p=0.683 Fig S2A) in their sensilla densities, nor did males and females (x̵_Male_=3414±1720 per mm^2^, x̵_Female_=2864±1128 per mm^2^; DFDen=38, F= 2.92, p=0.0956, Fig S2B). However, the interaction (activity*sex) was significant (DFDen=38.0, F=5.93, p=0.0197) with significantly higher sensilla densities on the antennae of nocturnal males compared to nocturnal females (x̵_Male_=3503±2145 per mm^2^, x̵_Female_=2395±1118 per mm^2^; Student’s t ratio= -3.17, p=0.003, Fig S2C).

The total sensilla counts included counts of three types of mechanoreceptors, nine types of chemoreceptors and one possible thermo/hygro receptor (Table 2). Overall, there were many more mechanoreceptors (in 1000s) than chemoreceptors (in 100s) on individual firefly antennae. *Phausis* had lower numbers (in 10s) for both sensilla types, but still ∼4 times more mechanoreceptors than chemoreceptors (Table 2). Potential thermo/hygroreceptors (X1) were identified in only a few individuals and, if present, in low numbers (N=1-22; Table S6).

### Testing predictions

We hypothesized that the differences in mating signals between diurnal and nocturnal fireflies would be reflected in their antennal sensilla counts and possibly in their sensilla densities. To test our predictions that diurnal species should have more chemoreceptors (including pheromone receptors) than nocturnal species, and that males should have more chemoreceptors than females, we conducted a comparative analysis of mechanoreceptors (no differences expected) and chemoreceptors (differences expected). We further predicted that diurnal males that rely on tracking a pheromone plume to conspecific females, would have more chemoreceptors than diurnal females.

#### Mechanoreceptor counts

Species accounted for 81.75% of the variation in mechanoreceptor counts across firefly antennae. Diurnal fireflies did not differ significantly in their mechanoreceptor counts from nocturnal fireflies (x̵_Diurnal_= 3503±993, x̵_Nocturnal_ = 1753 ± 1009; DFDen=5.0, F=4.88, p=0.078; Fig 8A), and males did not differ significantly from females (x̵_Male_ = 2586±1248, x̵_Female_ = 2420±1418; DFDen=33.0, F=0.21, p=0.279; Fig 8B). The interaction term (activity*sex) was not significant.

**Fig 8.**
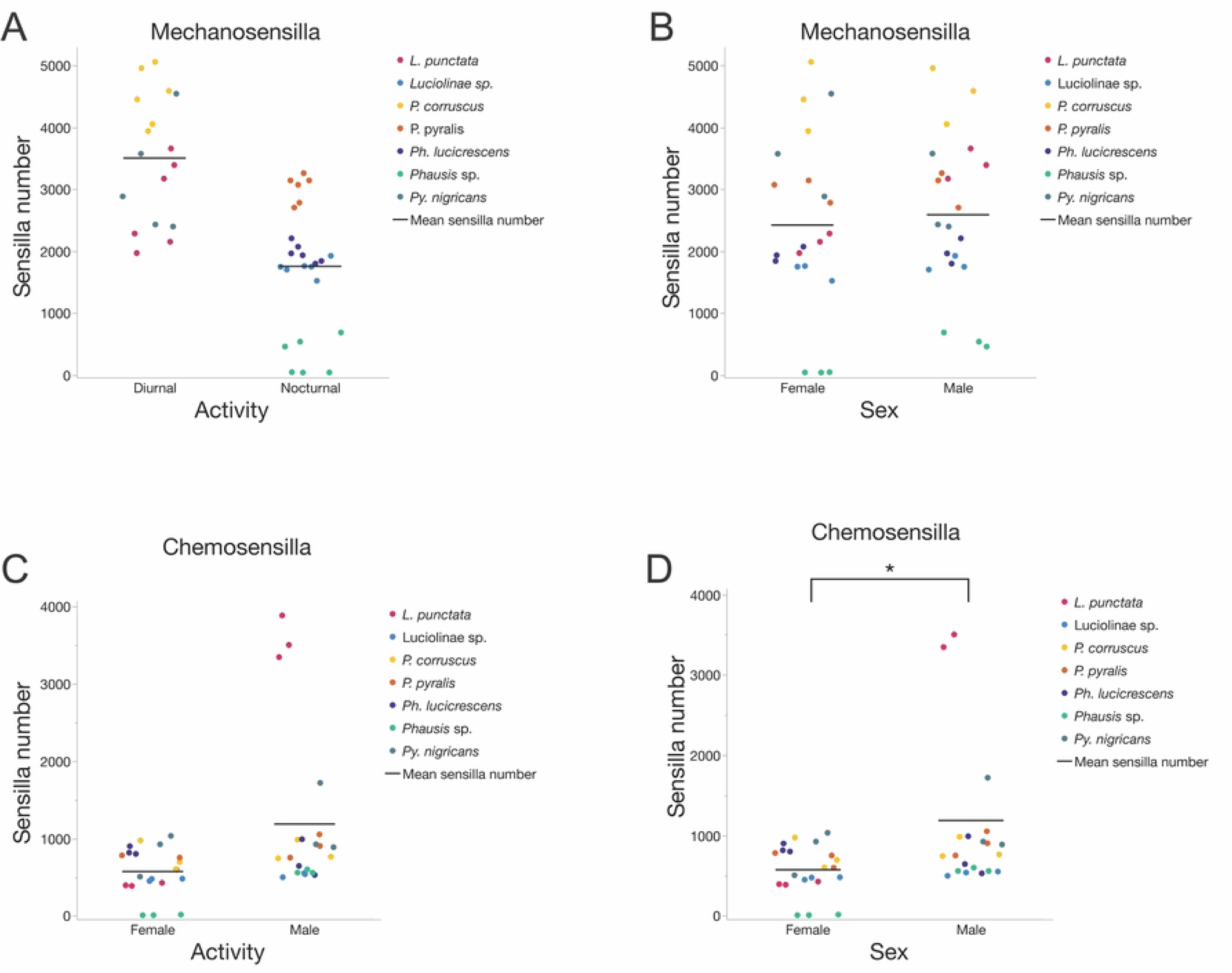
Mechanosensilla and Chemosensilla counts by activity and sex. (A) Mechanosensilla counts (both sexes) by activity time: there is no significant difference between diurnal and nocturnal species (x̵_Diurnal_= 3503±993, x̵_Nocturnal_ = 1753 ± 1009; DFDen=5.0, F=4.88, p=0.078); (B) Mechanosensilla counts (both activity times) by sex: there is no significant difference in mechanosensilla number between males and females (x̵_Male_ = 2586±1248, x̵_Female_ = 2420±1418; DFDen=33.0, F=0.21, p=0.279); (C). Chemosensilla counts (both sexes) by activity time: there is no significant difference between diurnal and nocturnal species (x̵_Diurnal_=1261±1114, x̵_Nocturnal_ = 595±282; DFDen=5.0, F=3.81, p=0.108); (D) Chemosensilla counts (both activity times) by sex: males have significantly more chemosensilla than females (x̵_Male_ = 1188±1041, x̵_Female_ =573±307; DFDen=33.0, F=15.66, p=0.0004).

#### Chemoreceptor counts

Species accounted for 32.27% of the variation in chemoreceptor counts across firefly antennae. Diurnal fireflies did not significantly differ in their chemoreceptors counts from nocturnal fireflies (x̵_Diurnal_=1261±1114, x̵_Nocturnal_ = 595±282; DFDen=5.0, F=3.81, p=0.108; Fig 8C), but males had significantly more chemoreceptors than females (x̵_Male_ = 1188±1041, x̵_Female_ =573±307; DFDen=33.0, F=15.66, p=0.0004; Fig 8D). The interaction term (activity*sex) was significant (DFDen=33.0, F=8.73, p=0.0057), with significantly more chemoreceptors in diurnal males (x̵_DMale_ = 1862±1326) compared to diurnal females (x̵_DFemale_ = 660±259; Student’s t-ratio= -4.57, p<0.0001, Fig S3A) and also compared to nocturnal males (x̵_NMale_ = 682±195; Student’s t-ratio= 3.08, p=0.0155; Fig S3B). In contrast, diurnal and nocturnal females (x̵_DFemale_ = 660±259, x̵_NFemale_= 507±334; Student’s t-ratio= 0.4, p=0.7; Fig S3C) did not significantly differ in their chemoreceptor counts, nor did nocturnal males and females (x̵_NMale_= 682±195, x̵_NFemale_= 507±334; Student’s t-ratio= -0.77, p=0.45; Fig S3D).

#### Sensilla counts, sensilla densities and antennal area

Both mechanoreceptor (R^2^=0.313, p=0.0001; Fig S4A) and chemoreceptor (R^2^=0.223, p=0.0016; Fig S4B) sensilla counts were positively correlated with antennal area, which means larger antennae had more sensilla of each type. This raises the question to what extent sensilla counts can be explained by antenna size, and/or whether there is also direct selection on sensilla numbers, e.g., due to activity time or sex of the respective firefly, resulting in higher sensilla densities. Overall, mechanoreceptor (R^2^= 0.549, p=0.0001; Fig S4C) and chemoreceptor (R^2^= 0.199, p=0.0031; Fig S4D) sensilla densities were negatively correlated with antennal area; this means that smaller antennae tended to have higher densities of both sensilla types.

#### Mechanoreceptor densities

Species accounted for 78.71% of the variation in mechanoreceptor densities. There was no significant difference between diurnal and nocturnal fireflies (x̵_Diurnal_= 2576/mm^2^ ± 827, x̵_Nocturnal_ = 2068/mm^2^ ± 1079, DFDen=5.0, F=0.477, p=0.52, Fig 9A) or between males and females in mechanoreceptor densities (x̵_Male_= 2265±1102 per mm^2^, x̵_Female_ = 2306±915 per mm^2^; DFDen=33.0, F=0.558, p=0.460; Fig 9B). However, the interaction term (activity*sex) was significant (DFDen=33.0, F=11.085, p=0.0021). This was due to significantly higher mechanoreceptor densities in diurnal females compared to diurnal males (x̵_DFemale_= 2887±571 per mm^2^, x̵_DMale_= 2265±954 per mm^2^; Student’s t-ratio=2.7, p=0.0109, Fig S5A) after species differences were taken into account as a random effect. In contrast, nocturnal females tended to have lower mechanoreceptor densities than nocturnal males (x̵_NFemale_ = 1870±896 per mm^2^, x̵_NMale_ = 2265±1243 per mm^2^; this trend was marginally significant: Student’s t-ratio= -1.97, p=0.057; Fig S5B). In comparison, diurnal and nocturnal females (x̵_DFemale_= 2887 ±571per mm^2^, x̵_NFemale_=1871±896per mm^2^; Student’s t-ratio=1.35, p=0.0.23; Fig S5C) and diurnal and nocturnal males (x̵_DMale_=2265± 954 per mm^2 x̵^_NMale_ =2265±1243 per mm^2^; Student’s t-ratio= - 0.00, p=0.999; Fig S5D) did not differ from each other.

**Fig 9:**
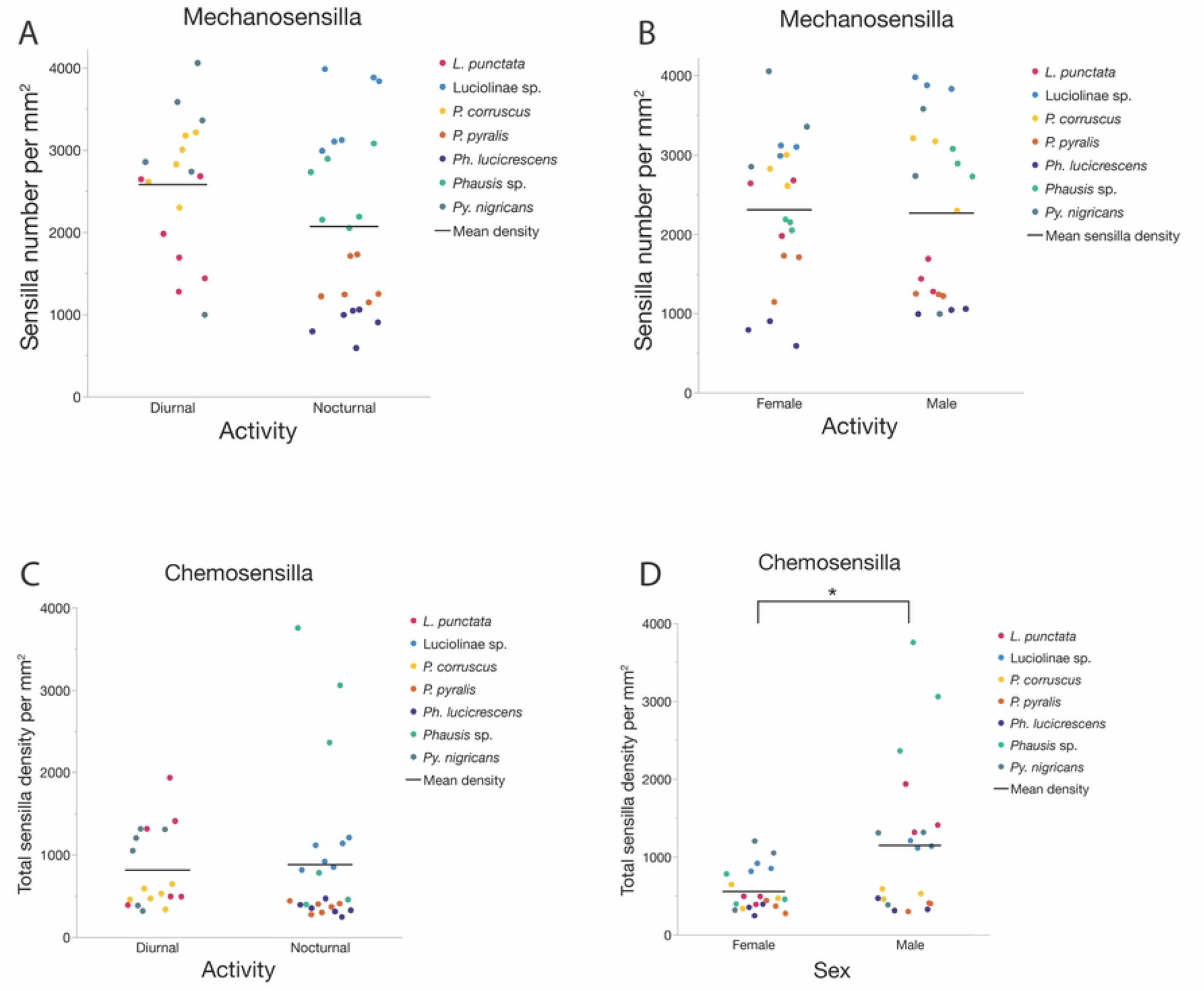
Mechanosensilla and Chemosensilla densities. (A) Mechanosensilla densities (both sexes) by activity time: there is no significant difference between diurnal and nocturnal species (x̵_Diurnal_= 2576/mm^2^ ± 827, x̵_Nocturnal_ = 2068/mm^2^ ± 1079, DFDen=5.0, F=0.477, p=0.52); (B) Mechanosensilla densities (both activity times) by sex: males have a significantly higher density of mechanosensilla than females (x̵_Male_ = 2265±1102 per mm^2^, x̵_Female_ = 2306±915 per mm^2^; DFDen=33.0, F=0.558, p=0.460); (C) Chemosensilla densities (both sexes) by activity time: there is no significant difference between diurnal and nocturnal species (x̵_Diurnal_= 813±486 per mm^2^, x̵_Nocturnal_ = 881±914 per mm^2^; DFDen=5.0, F=0.025, p=0.880); (D) Chemosensilla densities (both activity times) by sex: males have a significantly higher density of chemosensilla on their antennae than females (x̵_Male_=1147±952 per mm^2^, x̵_Female_=556±273 per mm^2^; DFDen=33.0, F=11.05, p=0.0022).

#### Chemoreceptor densities

Species accounted for 46.3% of the variation in chemoreceptor densities. Diurnal fireflies did not significantly differ in their chemoreceptor densities from nocturnal fireflies (x̵_Diurnal_= 813±486 per mm^2^, x̵_Nocturnal_ = 881±914 per mm^2^; DFDen=5.0, F=0.025, p=0.880; Fig 9C). Males had significantly higher chemoreceptor densities than females (x̵_Male_=1147±952 per mm^2^, x̵_Female_=556±273 per mm^2^; DFDen=33.0, F=11.05, p=0.0022; Fig 9D). The interaction term (activity*sex) was not significant.

### Sensilla Distribution on Firefly Antennae

To determine which portions of firefly antennae may be used for mechanoreception and/or chemoreception, we examined the distribution of sensilla numbers (mechanoreceptors, chemoreceptors, and each sensilla morphotype) across antennomeres. Mechanoreceptors were most abundant at the base of the antenna. C1 mechanosensilla occurred in high numbers on the scape, dropping to the lowest numbers on the pedicel (the second and smallest antennomere) in all species, except *Phausis* sp. (Fig S6G). C1 sensilla sharply increased in number from the pedicel to the third antennomere and occurred at similar numbers between antennomeres 4 and 10 for all species. Numbers further increased in *Photinus corruscus*, *L. punctata*, and *Photinus pyralis* on antennomere 11 (Fig S6-F). C2 sensilla were exclusively located on the scape and pedicel in all species, with about twice as many C2 sensilla found on the scape (Fig S6H-N). SC sensilla were absent in *Phausis* and in some sexes of other species (Fig S6O-T). SC sensilla were found on all 11 antennomeres but occurred in low numbers and were not evenly distributed. The antennomeres with the highest SC numbers differed between individuals of the same species and between species.

In contrast, chemoreceptor sensilla were entirely absent from the scape and pedicel. They tended to be most abundant in the middle of the firefly antennae. B1 sensilla increased in number between antennomeres 3 to 6 and then decreased from antennomere 6 to 10 in *Photinus corruscus* and *P. pyralis*. In contrast, the distribution of B1 sensilla in *Pyropyga nigricans* were overall consistent across antennomeres 3-8 (Fig S7A-C). The number of B2 sensilla also increased between antennomeres 3 and 6 and then decreased from antennomere 6 to 11 (Fig S7D-E). B3 sensilla numbers varied between antennomeres 3 to 8 and gradually decreased in number in each subsequent antennomere (Fig S7F-H). B7 sensilla were found on antennomere 3 in all species except *P. pyralis*. B7 numbers varied between antennomeres 3 to 10 with a sharp increase on antennomere 11 in all species, except *Photinus pyralis* (B7 decreased on antennomere 11; Fig S7I-N). B10 sensilla (*Lucidota punctata*) were most abundant between antennomeres 4 and 6 (Fig S7A), slowly decreasing in number towards the distal antennomere; males have about twice as many B10 on each antennomere (peak average of 99 sensilla on antennomere 4) compared to their females (peak average of 56 on antennomere 6). B11 sensilla (*Lucidota punctata*) were rare on female antennae with most (average 9 sensilla) located on the last antennomere; in contrast males consistently averaged between 300 and 350 B11 sensilla on antennomeres 3-11 (Fig S8B.). B12 sensilla (Luciolinae sp.) were present in low numbers on antennomeres 3-10 of male antennae, with the highest number (average 2.7 sensilla) on antennomeres 3 and 4 and the lowest number (average 0.33 sensilla) on antennomere 5 (Fig S8C). B12 sensilla were absent in females. B13 sensilla (Luciolinae sp.) were present on antennomeres 3 to 11 on male antennae, with the highest number on antennomere 5 (average 81 sensilla), then slowly declining in numbers to antennomere 11 (average 17 sensilla; Fig S8D). In parallel to males, B13 sensilla of females were present on antennomeres 3 to 11, with the highest number on antennomere 5 (average 71 sensilla), slowly declining to antennomere 11 (average 14 sensilla; Fig S8D). T1 sensilla (*Phausis* sp.) were present on the 2nd antennomere (of 3) on female antennae (3-10 sensilla each; Fig S8E). *Phausis* males had T1 sensilla on antennomeres 3-11, and they were more or less evenly distributed (ranging on average 55-70 sensilla on each antennomere) between antennomeres 3 to 11 (Fig S8E). X1 sensilla were only found in three species: *Photinus corruscus, Pyropyga nigricans*, and *Photuris lucicrescens*, and only on individual fireflies. In *Pyropyga. nigricans* and *Photinus corruscus,* X1 sensilla were present exclusively on the ventral side of the antenna, and on both dorsal and ventral sides in a single female of *Photuris lucicrescens*.

## Discussion

We report here a total of 14 sensilla morphotypes across seven species of Lampyridae. Twelve of these morphotypes are new for Lampyridae. The two other morphotypes were previously reported for *Luciola cruciata* [23], along with five morphotypes not found in our study species. These five morphotypes include different variants of sensilla campaniform, sensilla basiconica, sensilla trichodea, and two additional sensilla categories: capitular sensilla and gemmiform sensilla, the latter being a new type for beetles. This results in a total of 19 recorded sensilla morphotypes for 8 species of Lampyridae and suggests that with the further addition of firefly taxa new morphotypes await to be discovered. The majority of these 19 morphotypes are also found in other beetle groups, except for gemmiform sensilla in *Luciola cruciata* and B10 and B11 sensilla of *Lucidota punctata*, which are reported and described for the first time here. The B10 and B11 sensilla of *L. punctata* are unique in their wide-collared base and the shape of their peg (Fig 4D, E).

In contrast to the more diverse chemoreceptors, there are only three types of mechanoreceptors on firefly antennae (two variants of sensilla chaetica and one sensilla campaniform). These tend to occur in much greater numbers (in 1000s) than chemoreceptors (in 100s; Table 2), and the three mechanoreceptor types are widely distributed across fireflies. For example, both males and females of all eight firefly species studied so far have sensilla chaetica types C1 and C2 on their antennae. C1 mechanosensilla are present on all 11 antennal segments (Fig S6A-F). They tend to occur in high numbers on the scape, drop in numbers on the pedicel (the smallest antennomere in all species, with the exception of *Phausis* sp. males and females), then sharply increase in numbers on antennomere 3, and remain at relatively high numbers on the other antennomeres. This places C1 sensilla in a good position to process mechanical stimuli from the environment along the entire length of the antennae. C1 sensilla are also consistently found in in closely related Elateridae species, in more distantly related beetles (e.g., Chrysomelidae and Carabidae), and across Insecta (e.g., Blattodea, Hemiptera) in general [15, 55–59]. Like in other elaterid groups [15, 60], the C2 sensilla of fireflies were exclusively found on the first two antennomeres (scape and pedicel), where the major muscles for antennal movement are located [2], supporting their function in antennal proprioception [6, 61, 62].

The third type of mechanoreceptor sensilla, sensilla campaniform (SC sensilla), is relatively rare in fireflies. Six of the eight firefly species studied to date (except *Phausis* sp. and *L. cruciata*) have SC sensilla (Table 4), but they were absent in *Lucidota punctata* males and *Photuris lucicrescens* females. This could be possibly due to the overall low numbers of SC sensilla on firefly antennae. SC sensilla were most prevalent in *P. pyralis* and *Py. nigricans* (Table S2). SC sensilla are commonly found on insect legs and insect wings and typically function as strain sensors within the exoskeleton [63]. When stimulated, SC sensilla have been found to trigger muscle contraction for stability or propulsion [64], however, whether SC sensilla on insect antennae could stimulate the antennal support muscles in the scape and pedicel remains to be tested.

With nine sensilla morphotypes, chemoreceptors are much more diverse than mechanoreceptors on firefly antennae, and their distribution across species is unexpectedly variable (Table 4). Chemoreceptor sensilla were absent from the scape and pedicel of firefly antennae, but present on all other antennomeres (3-11), with the exception of *Phausis* sp. females. They tend to be most numerous in the middle and/or the distal end of the antenna (Figs S6 and S7). The chemoreceptor sensilla type shared by most species (except *Phausis* sp.) were B7 sensilla, which are found in both male and female fireflies, but in very low numbers (Table S4). Their function in fireflies is unknown, but B7-like sensilla in Cerambycidae and Curculionidae are hypothesized as thermo/hygro receptor sensilla, based on their internal cell structure [65, 66].

The other chemoreceptor sensilla types were shared by three firefly species (B1, B3), two firefly species (B2 sensilla), or they were unique for a single species (B10, B11, B12, B13, T1). Sensilla basiconica B1, B2, B3, and B7 are also present in Elateridae [15], in Chrysomelidae [67], and Cerambycidae [68]. Most sensilla basiconica are hypothesized to be olfactory sensilla based on the presence of pores [69]. B1 and B2 like sensilla are hypothesized as olfactory sensilla in Chrysomelidae [70], B3 sensilla have been hypothesized as olfactory sensilla in both Cerambycidae: *Xylotrechus* and Elateridae: *Tetrigus* [71, 72], and B7 sensilla are hypothesized as either olfactory or thermoreceptive sensilla in Cerambycidae: *Aromia* [68]. Sensilla trichodea (T1) detect sex pheromones in pine weevils (*Hylobius abietu*, Curculionidae [20]) and are the only chemosensilla in *Phausis* fireflies. Whether they are used by *Phausis* to sense pheromones [31] remains to be tested. Sensilla coeloconica sensilla (X1), while lacking visible pores, should also be considered as potential chemosensilla for either olfactory or contact chemicals. The grove on the distal end of these sensilla may conceal pores or openings in the sensilla wall, which would allow chemicals to enter and stimulate nervous cells. X1 sensilla have been identified as temperature sensors in *Siagona europaea* (Carabidae [22]), further supporting the possibility of an opening in the sensilla wall that would allow fluid to pass through the sensilla wall. They are present in low numbers on the antenna of *Photinus corruscus*, *Pyropyga nigricans*, and *Photuris lucicrescens* and it remains to be tested whether the X1 neurons in these species respond to temperature or humidity stimuli. Interestingly, X1 sensilla are restricted to the ventral antennal surface of *Photinus corruscus* and *Pyropyga nigricans* and the functional significance of this sensilla distribution remains unknown. Sensilla coeloconica are also hypothesized as thermoreceptors in Diptera [73]. Cone sensilla resembling the ones found in fireflies, were observed in the Elateridae species *Agriotes lineatus* [15], but only in a single specimen. The cuticular pores observed in this study closely resemble the glandular pores associated with C1 sensilla in *Agriotes* [15]. Similar pores were also observed in *Drilus mauritanicus* and *Drilus flavescens* [60, 74] and labeled as perforated plates. The function of these large pores remains unknown but they are hypothesized to be associated with glandular secretions [74–77].

### Evolution of sensilla types

Given our taxon sampling of eight taxa in six genera and their phylogenetic relationships [51], it appears that mechanoreceptor sensilla types are relatively conserved and shared by all (C1) or almost all (C2, SC) firefly species studied to date. In contrast, their chemoreceptor sensilla types are much more diverse and only a single sensilla type (B7) is present in all species and thus seems to be inherited from the common ancestor of our study species (Fig 6). The spotty distribution of the other chemoreceptor sensilla types across firefly genera suggests a high evolutionary lability of chemoreceptor sensilla types among the six firefly genera in our study. However, these genera are relatively distantly related and belong to three different lampyrid subfamilies. On the other hand, the two *Photinus* species share many of the same sensilla types, despite their different (diurnal/nocturnal) activity times. To further investigate this seeming evolutionary lability of chemoreceptor sensilla types across genera and the conservation of sensilla types within a genus, more species in each genus need to be studied, as well as additional genera that are closely related to our study taxa [51].

### Optimizing sensitivity

Antennal sensitivity to relevant environmental stimuli can be increased through additional sensilla, thus increasing the probability of capturing physical or chemical stimuli. This can be achieved by increasing antenna size (surface area) or by packing sensilla more tightly (density). In the present study diurnal fireflies had significantly more sensilla on their antennae than nocturnal fireflies, and thus a higher sensitivity for environmental stimuli. Similarly, males had significantly more sensilla on their antennae than females. Neither diurnal and nocturnal fireflies, nor males and females differed in overall sensilla densities. Thus, in fireflies overall antennal sensitivity (sensilla numbers) seems to be mainly increased by increasing the antennal surface area. Larger antennae also have the advantage of sampling a larger airspace, which increases the probability of detecting the pheromones of a conspecific female [26]. Not surprisingly, firefly males - as the actively searching sex - have relatively larger antennae (relative to their body size) and significantly higher total sensilla counts compared to their females. The evolution of larger firefly antennae as an improved sampling area seems to be supported by the negative correlation between antennal area and sensilla density: as firefly antennae increase in size, sensilla numbers do not increase in a 1:1 relationship with antennal area, resulting in a lower density. However, there may be other limiting factors for sensilla density on firefly antennae, such as limited space in the antennomeres that supports a limited supply system for the receptor neurons in each sensillum [6].

Larger body size in insects supports larger antennae [78, 79], including in fireflies [35]. Not surprisingly, across our seven species, body size (pronotum length) had a significant effect on antennal area, with species accounting for almost all (95.06%) of the variation in body size. The body size of fireflies in our species sample did not differ significantly by sex or activity, thus body size did not influence our statistical analysis of sex and activity effects on sensilla counts or densities. This means that the significantly larger antennal areas of firefly males compared to females cannot be explained by a difference in body size. Instead, the evidence points towards direct selection on male antenna size, most likely sexual selection resulting in increased antennal sensitivity of male antenna to chemical stimuli. This is supported by significantly larger antennal areas of diurnal males compared to diurnal females.

In contrast, diurnal and nocturnal fireflies in our study overlapped greatly in their antennal areas, and did not differ significantly from each other. We suspect that this is likely due to our relatively small species sample. A previous large-scale analysis of eye and antenna sizes in 101 firefly taxa showed that the males of the 26 diurnal firefly taxa tended to have significantly longer antennae (and smaller eyes) than the males of the 75 nocturnal taxa [35] with activity time (and type of mating signals) accounting for 13% of the observed variation in male antenna size, while phylogenetic relatedness (genus) accounted for 63% [35]. Similarly, in a phylogeny-based analysis of 43 North American firefly species, antenna length (phylogenetic signal lambda=0.859) was significantly correlated with body size (pronotum length, p<0.0001) and activity time (mating signal: p=0.037). Combined with the results from the present study, it appears that the evolution of a larger antenna size is an important factor in increasing sensilla counts, and thus antennal sensitivity, in diurnal fireflies.

Interestingly, when fireflies change their activity time from nocturnal to diurnal over evolutionary time (and switch their mating signals from bioluminescent signals to the exclusive use of pheromones), the selection on antenna size seems to be relatively weak, resulting in a relatively slow increase in antenna size compared to the reduction in eye size [35]. For this reason, Stanger-Hall et al. [35] suggested that the first response to selection for improved pheromone detection may occur through sensilla numbers instead, resulting in an increased sensilla density on relatively short antennae. An analysis of our data for *Photinus pyralis* (N=6) and *P. corruscus* (N=6) shows that this is indeed the case. The diurnal firefly *P. corruscus*, which belongs to a clade of fireflies (formerly *Ellychnia* [47]) that split from their common ancestor with nocturnal *Photinus* fireflies between 6 and 25 mya [80, 81] has significantly shorter antennae (scaled with body size: Wilcoxon rank-sum test: S=56, Z=2.64, p=0.008) than the nocturnal *P. pyralis*, but it has significantly more sensilla (Wilcoxon rank-sum test: S=21, Z=-2.80, p=0.005) and a significantly higher sensilla density (Wilcoxon rank-sum: S=21, Z=-2.8, p=0.005). This is due to significantly higher mechanosensilla numbers (x̵ = 4506±456 versus x̵ = 3015±222.5, Wilcoxon rank sum N=6,6, S=21, Z=-2.8, P=0.005) and densities (x̵= 2851±352 per mm^2^ versus x̵= 1381±263 per mm^2^; (Wilcoxon rank sum N=6,6, S=21, Z=-2.8, P=0.005). Interestingly, the total chemosensilla counts of *P. corruscus* and *P. pyralis* did not differ (N=6,6, x̵= 794±154 versus x̵ = 806±155; Wilcoxon rank sum N=6,6, S=41, Z=0.24, P=0.81), but *P. corruscus* (with its shorter antennae) has a significantly higher chemosensilla density (x̵= 504±109 per mm^2^ versus x̵= 365±65 per mm^2^; Wilcoxon rank sum N=6,6, S=25, Z=-2.16, P=0.031) as predicted by Stanger-Hall et al. [35]. Given the similar chemoreceptor numbers for *P. corruscus* and *P. pyralis*, this raises the question whether (or to what degree) *P. pyralis* may still utilize pheromones during mate search.

### Testing predictions

We hypothesized that the differences in mating signals between diurnal and nocturnal fireflies will be reflected in their antennal sensilla counts and possibly in their sensilla densities. Given the importance of pheromones for the mate search of diurnal firefly species, we predicted (1) more chemoreceptors (including pheromone receptors) in diurnal species compared to the bioluminescent nocturnal species, and (2) more chemoreceptors in males compared to females, and specifically that diurnal males, which are tracking pheromone plumes to females, would have more chemoreceptors than their females. In comparison, we predicted no differences in mechanoreceptor counts and densities between diurnal and nocturnal fireflies and/or between males and females.

#### Mechanoreceptors

Species accounted for most (81.75%) of the variation in mechanoreceptor counts across firefly antennae, reflecting large species differences in sensitivity to mechanical stimuli, possibly due to different microhabitats. Diurnal and nocturnal fireflies did not differ in their mechanoreceptor counts, nor did males and females. There was also no significant interaction between sex and activity in mechanoreceptor counts. This finding was not unexpected, since both diurnal and nocturnal fireflies (and both sexes) need mechanoreceptors to navigate through their physical environment during mate search, to deposit fertilized eggs, and from and to their daily resting places. Similarly, species accounted for most (78.71%) of the variation in mechanoreceptor densities. There was no significant difference between diurnal and nocturnal fireflies or between males and females, however the interaction (sex*activity) was significant with significantly higher mechanoreceptor densities in diurnal females compared to diurnal males. This is likely due to the significantly smaller antennal areas of diurnal females compared to diurnal males, resulting in a denser packing of similar mechanosensilla numbers on a smaller area.

#### Chemoreceptors

Species accounted for a relatively small part (32.27%) of the variation in chemoreceptor counts across firefly antennae. Overall, the diurnal fireflies did not significantly differ in their chemoreceptors counts from the nocturnal fireflies in our study, but males had significantly more chemoreceptors than females and the interaction term (activity*sex) was also significant. As predicted, diurnal males had significantly more chemoreceptors than nocturnal males. Diurnal males also had significantly more chemoreceptors than their females, who release pheromones, but have not been documented to respond to them (e.g. to pheromones of other females). The females of diurnal and nocturnal fireflies did not differ significantly in their chemoreceptor counts, nor did nocturnal females and their males, who rely much less (if at all) on pheromones than the diurnal males. Species accounted for 46.3% of the variation in chemoreceptor densities. Overall, diurnal fireflies did not significantly differ from nocturnal fireflies, but males had significantly higher chemoreceptor densities than females. Since males also have significantly larger antennal areas than females, these results suggest that sexual selection in firefly males can act through both, a significantly larger antennal area and/or a higher density of chemoreceptors to optimize the sensitivity of male antennae to chemical stimuli. For example, *Phausis* and Luciolinae sp. males have relatively small antennae with high densities of chemoreceptors, and *Lucidota* males have both large antennae and a high chemoreceptor density (Tables 2, 3).

#### Identification of pheromone sensilla candidates in fireflies

Diurnal firefly species had three to five different chemoreceptor sensilla types on their antennae, while nocturnal species had one or three (two types were reported for the nocturnal species *Luciola cruciata* [23]). This trend directly reflects the importance of chemoreceptors for diurnal species and is further supported by a significantly higher number of chemoreceptors in diurnal males compared to nocturnal males (Fig S3B). However, the absence of a specific chemoreceptor sensilla morphotype found exclusively in diurnal firefly species suggests that pheromone sensilla may not be unique for diurnal species. If pheromone sensilla are not unique to diurnal species, the most likely pheromone sensilla candidate is B7, a grooved-peg sensilla type that is shared by 6 of the 7 species in our study (except *Phausis* sp.). Grooves on B7 pegs may conceal openings in the sensilla wall that would allow chemical stimuli to pass through, indicating them a possible candidate for pheromone receptors. However, B7 sensilla were found in relatively low numbers compared to other chemoreceptors and with inconsistent differences between males and females (Table S4). Furthermore, B7 sensilla may be used as thermoreceptors and/or hygroreceptors, as is known for other grooved peg sensilla [68, 82–84]. If this were indeed the case for fireflies, supported by the relatively low and similar numbers in males and females, B7 would serve an essential function for the survival of these soft-bodied and thus easily dehydrated beetles. However, *L. cruciata* fireflies are using their unique capitular sensilla for that purpose [23], which raises the question what their B7 sensilla are used for. If *Phausis* is able to sense temperature or humidity with antennal sensilla, it would be using T1 sensilla, its only non-mechanoreceptor sensilla type. All these possibilities await future functional testing.

The absence of universal “firefly pheromone sensilla” for diurnal species, or for both diurnal and nocturnal species and the limited overlap of the different chemoreceptor sensilla types between genera, suggests that pheromone sensilla, if present, may be genus- or even species-specific.

Therefore, we applied our predictions for pheromone sensilla candidates to the most abundant chemical sensilla morphotypes found in each species (Table S4) to propose specific sensilla types for future testing. B1 sensilla are the most abundant chemoreceptors in diurnal *Photinus. corruscus* fireflies and males have almost twice as many B1 than their females. Males have on average ∼50 B1 sensilla on most of their antennomeres (antennomeres 4-10), but B1 numbers peak to an average of ∼75 sensilla on the distal (11th) antennomere (Fig S7), enabling them to sample a large airspace with their two antennae and increasing their chances to catch molecules from the female pheromone plume. Most importantly, basiconica sensilla that look most like the B1 sensilla in our study were identified through electro-antennograms as pheromone sensilla in diurnal *P. corruscus* [21]. However, pores, if present, are not visible in our SEM images; this could possibly be due to the gold-coating obstructing small pores [85]. Based on their numbers, B2 and B3 sensilla could be two other pheromone sensilla candidates, but in both cases, females have more B2 and B3 sensilla than males. Furthermore, sensilla similar in morphology to B3 sensilla were unresponsive to *P. corruscus* sex pheromones [21], confirming that B3 sensilla are not used for the detection of sex pheromones in this species.

The nocturnal *P. pyralis,* a closely related congener of *P. corruscus* in this study, also has high numbers of B1 sensilla, and males have ∼ 30% more B1 sensilla than their females (Table S4), suggesting that both *Photinus* species may use B1 sensilla as pheromone sensilla. However, in contrast to *P. corruscus*, in *P. pyralis* males most B1 sensilla (>50/antennomere) are located in the center of their antennae (antennomere 4-9; Fig S7C). The next closest relative to these two *Photinus* taxa in our study is *Pyropyga nigricans,* a diurnal firefly species. It also has B1 sensilla, possibly inherited from a common ancestor with *Photinus* fireflies (Fig 6), however at very low numbers (∼5/antennomere) and, contradictory to our pheromone sensilla predictions, females have twice as many B1 sensilla than males. This makes B1 sensilla an unlikely pheromone sensilla candidate for *Pyropyga*. In contrast, pored B3 sensilla are present in great numbers (>100/antennomere on antennomere 3-11; Fig S7F-H), and *Py. nigricans* males have ∼1.5 times more B3 sensilla on their antennae than their females (Table S4), suggesting B3 as pheromone sensilla candidate for *Py. nigricans*.

The species-specific B10 and B11 sensilla make good candidates as pheromone sensors for the diurnal species *L. punctata*. Males have ∼1.5 times more B10 sensilla and ∼100 times more B11 sensilla than their conspecific females. In addition, B10 sensilla have obvious pores. B10 sensilla are found in the highest numbers between antennomere three and eight in both males and females, while B11 sensilla are found in equal numbers across antennomeres three to eleven in both males and females, but females have a slight increase in B11 sensilla on their distal (11th) antennomere.

In the nocturnal Luciolinae species, the species-specific B12 and B13 sensilla are both candidates for pheromone receptors. B12 were found in low numbers (13±3) exclusively on the antennae of males between the pedicel and antennomere 9. B13 sensilla occurred in much higher numbers in males (488±29 per antenna), but were also present in just slightly lower numbers (442±29) on the antennae of females, with the highest numbers on antennomere 5 in both sexes (Fig S8C, D: distribution B12 and B13). Based on our prediction of sexual dimorphism this makes B12 a slightly more likely pheromone sensilla candidate, however the very low numbers would suggest that pheromones do not play an important role for the Luciolinae species. Alternatively, based on total sensilla numbers (antennal sensitivity), B13 would be a good pheromone candidate, if pheromones are indeed used by *Luciolinae* sp., which remains to be studied.

The only chemoreceptor sensilla found in nocturnal *Phausis* sp. were species-specific T1 sensilla. These were found in much greater numbers (∼50x) in *Phausis* sp. males compared to females, making T1 sensilla the likely (and only) candidate for pheromone detection in this species [31], and possibly also in the sister genus of *Phausis*, *Lamprohiza,* for which the dual use of bioluminescence and pheromones has just been confirmed [40]. In nocturnal *Ph. lucicrescens*, B2 sensilla are the most prevalent chemoreceptor type, but females have ∼100 times more B2 sensilla than their males. This either suggests that nocturnal *Ph. lucicrescens* rely exclusively on bioluminescence and therefore do not have any pheromone sensilla, or that B2 sensilla may serve more than one function. The predatory females of *Ph. lucicrescens* may use their B2 sensilla to recognize prey species, specifically *Photinus* fireflies, which sequester lucibufagins [86, 87], potent defense chemicals against ants and vertebrate predators [88]. The greater abundance of B2 sensilla in *Ph. lucicrescens* females compared to their conspecific males may aid these predatory females to identify captured *Photinus* prey (via cuticular hydrocarbons or other low-volatile chemicals). Whether this is indeed the case, and whether the same sensilla are used by males to detect female pheromones (if any), remains to be tested.

### Gustation during antennation behavior

The intensive antennation behavior that precedes mating in fireflies, led us to hypothesize the sampling of chemical cues by gustatory (contact) chemoreceptors to verify a conspecific mate, immediately before mating. Based on this reciprocal behavior we predicted no sex differences in sensilla that pick up on these cues. We presently do not know (1) to what extent individual firefly species use contact chemicals during mating (or whether antennation represents a purely physical stimulus in preparation for mating), and (2) whether the same sensilla could function both as pheromone sensilla (olfactory receptors stimulated by high-volatile chemicals) and gustatory sensilla (taste receptor cells stimulated by low-volatile contact chemicals on the cuticle or released during mating), or whether different sensilla types process these different stimuli. For example, the same sensilla type could be used to process both stimuli with specialized olfactory and gustatory receptors on their sensory neurons at the base of the sensillum [89]. In this case, we would predict a less pronounced sexual dimorphism in candidate sensilla numbers compared to sensilla that function exclusively as pheromone sensilla. Alternatively, if different sensilla types are used, we would predict one chemosensilla type with a pronounced sexual dimorphism (olfactory pheromone sensilla) and a second chemosensilla type (gustatory contact chemical sensilla) without sexual dimorphism, or even skewed towards females, because they incur a larger cost for a mating mistake. Based on these criteria, we propose the following candidates for future functional testing as gustatory sensilla: For separate (specialized) gustatory sensilla we propose B2 or B3 for *Photinus corruscus*, B3 for *P. pyralis*, B1 for *Py. nigricans*, B10 for *L. punctata*, and B13 for Luciolinae sp. If no pheromones are used by *Photuris lucicrescens*, B2 would test as a gustatory sensilla, with a sexual dimorphism based on predatory behavior of females. However, if pheromones are used by *Photuris lucicrescens*, B2 sensilla represent a potentially mixed sensilla type that processes both female pheromones and gustatory prey signatures in this species. For *Phausis* sp. with a single chemosensilla type, T1 sensilla would both function as pheromone sensilla, as well as gustatory sensilla used by males during their antennation behavior [31].

Possible other uses of chemoreceptors in fireflies include the localization of milkweed for nectaring. Fireflies from three genera (*Photinus*, *Pyropyga*, *Photuris*) have been observed to collect nectar from milkweed, choosing only the freshest aromatic flowers, and two additional genera (*Lucidota*, *Pyractomena*) have been seen on milkweed [90]. In addition, *P. corruscus* has been observed feeding on the floral nectaries of Norway maples [91]. Both males and females show this behavior, therefore no sex differences in the respective chemoreceptors would be expected.

As an important next step in understanding how fireflies perceive the world through their antennae, and how this influences their behavior, all of the pheromone and gustatory sensilla candidates need to be tested in functional studies. One key question in this context is the extent to which the pheromone sensilla candidates remain functional in nocturnal firefly species. Our morphological data suggest that B1 sensilla are pheromone sensors in both *Photinus* species: *P. pyralis*, a nocturnal bioluminescent species, and *P. corruscus*, which split from nocturnal *Photinus* ∼6-25 mya [80, 81] and returned to diurnal activity with the exclusive use of pheromones for mate search [34, 37]. Other repeated losses of bioluminescence and reversals to mate search exclusively with pheromones during firefly evolutionary history [36, 37], suggest that pheromone sensilla may remain functional at least in some nocturnal species, especially in those clades with recent reversals to diurnal activity (e.g., *Photinus*), facilitating the switch from nocturnal to diurnal activity. If functional in nocturnal species, pheromones would increase mating opportunities, because they would attract and direct males towards the bioluminescent display sites of females. Once they are close enough, they can use visual cues to locate and/or identify conspecific females over shorter distances.

Pheromones are especially important in environments that are not conducive to visually finding females. For example, considering their antennal areas, the males of diurnal *Lucidota* and of nocturnal *Phausis* both have relatively high chemosensilla (but not mechanosensilla) densities on their antennae. *Lucidota punctata* (5-6mm body length) and *Phausis reticulata* (6-9mm) are both relatively small fireflies and the males of both species search for conspecific females in the low vegetation on the forest floor, navigating around leaves to find their females [92]. The diurnal *L. punctata* males have relatively large antennae, combined with a relatively high chemosensilla density to locate their tiny females in shady forests. The males of nocturnal *Phausis reticulata* (blue ghosts) fly at night and glow while trying to locate their tiny flightless females. *Phausis* females emit a weak glow (visible within 10 feet, but almost invisible when moonlight is reflected from wet leaves [92]), and *Phausis* males seem to use female pheromones to get close to females. If the male cannot locate the female by her weak glow, males may use their own glow as “spotlight” to locate the female [31].

### Why are chemosensilla so diverse across fireflies?

The individual firefly species in our study had between one to four different chemoreceptor sensilla types on their antennae. The number of types seem to be correlated with diurnal or nocturnal activity, however, a puzzling insight from our study is the diversity of antennal sensilla types in fireflies, and how little overlap there is between genera. The adult fireflies in our study tend to survive for only a few weeks (except for the winter firefly *P. corruscus* which may live several months [92]) for the sole purpose of mating, and diurnal and nocturnal species face very similar challenges: prevent dehydration, locate a conspecific mate, mate, and in the case of females, find a suitable egg deposition site. So why are their antennal chemosensilla so different? We identified three different pheromone sensilla candidates in our three diurnal firefly species (B1, B3, B10/B11), which raises the question of how it is possible that these different sensilla morphotypes converged on an apparently similar function?

In *Drosophila* trichoid sensilla are required for pheromone detection, while their basiconic sensilla mostly detect food-derived odors [46]. Similarly, trichoid sensilla respond to female pheromones in click beetles (Elateridae; 61) and in Asian longhorned beetles (Cerambycidae; 94), while placoid sensilla detect pheromones in Japanese beetles (Scarabaeidae; 95). In fireflies (Lampyridae) we identified trichoid sensilla as pheromone sensilla candidates for nocturnal *Phausis*, however, trichoid sensilla are absent in all other firefly species studied to date. Instead, sensilla basiconica are used for pheromone detection in the firefly *Photinus corruscus* [21]. *P. corruscus* females emit (1*S*)-*exo*-3-hydroxycamphor (hydroxycamphor), which in single sensillum recordings elicited a neuronal response from a pheromone-sensitive olfactory sensory neuron in a basiconica sensillum on the male antenna [21]. We identified this sensillum as sensilla basiconica type 1 (B1). A possible model for how pheromone detection can switch between major sensilla types, is the Asian longhorned beetle *Anoplophora glabripennis* (Cerambycidae), whose pheromone consists of two components [94]. Trichoid sensilla are the pheromone sensors that respond to both components, but basiconica sensilla respond to one of the two components (along with plant compounds that enhance male attraction). Even though there is no pheromone-specific information relayed by these basiconica sensilla, this observation suggests a possible mechanism for sensilla type switching as pheromone compounds diverge in closely related species. Similarly, in the moth *Ostrinia nubilalis* (Crambidae) pheromone receptors that respond to different pheromone compounds are split between two olfactory sensory neurons in two sensillum subtypes, which is thought to reflect an ongoing evolution of this sensillum type as two *O. nubilalis* strains diverge [96]. A similar mechanism could account for the diversification of B-sensilla in fireflies (Lampyridae) and the diversity of pheromone sensilla candidates across diurnal firefly species. Sensilla type switching may not be limited to fireflies, but reflect the evolutionary dynamics of pheromone signals and their receptors (sensory neurons) in insects in general; evolutionary studies with a broad taxon sampling across closely related species in different genera will be required to test this.

## Conclusions

This study presents the most comprehensive description of the antennal sensilla of Lampyridae to date. We identified 12 new firefly sensilla morphotypes, for a total of 19 morphotypes documented now for 8 species in 6 genera. We documented these sensilla types for both males and females of our 7 study species. Mechanoreceptor sensilla were the most abundant sensilla on firefly antennae, but chemoreceptor sensilla were unexpectedly diverse. Fireflies mainly increase the sensitivity of their antennae to environmental stimuli by increasing their antennal area, and this translates into more sensilla, as documented here. In addition, male fireflies may increase their sensilla densities. As predicted, diurnal and nocturnal fireflies did not differ in their mechanoreceptor counts or densities, nor did males and females. But males had significantly higher chemosensilla counts and densities than females, underlining the importance of males and their chemoreceptors (including pheromone sensilla) for locating a conspecific female by tracking her pheromone plume. Reflecting their exclusive reliance on pheromones during mate search, diurnal males had significantly more chemosensilla (but not higher densities) than nocturnal males, while diurnal and nocturnal females did not differ. An increase in chemoreceptor sensilla density may be utilized in male fireflies with relatively small antennae (e.g., *Phausis*), and/or by males with large antennae (e.g., *Lucidota*) to further optimize antennal sensitivity, especially when females are difficult to locate in their environment.

We did not identify a “universal pheromone sensilla” candidate for diurnal (and/or nocturnal) fireflies, but we used our predictions for pheromone sensilla to propose candidate sensilla types for the different species based on their respective sensilla numbers and morphology. These pheromone sensilla candidates will facilitate functional testing in future studies. Similarly, we identified potential candidates for gustatory recognition (if any) of conspecifics during antennation, which could be utilized by different species. It is currently not known whether gustatory chemicals are sampled during antennation or whether this behavior serves as purely physical stimulation in preparation for mating. Olfactory and/or gustatory chemoreceptors may also be used for nectaring by both male and female fireflies. While our study was limited to firefly species with filiform antennae, our study revealed an unexpected diversity of sensilla types in fireflies. Most of the antennal forms known for beetles (except clavate and plumose) occur within the firefly family, Lampyridae [2, 29]. We predict that future studies will uncover an even greater sensilla diversity across Lampyridae and inform us whether antennal sensilla cover the entire surface of more complex firefly antennae or whether these enhanced 3-dimensional structures serve other functions, e.g., the direction of airflow across the antennae [97]. We propose both morphological and functional studies of antennal sensilla with a broad taxon sampling of closely related species across genera to illuminate the dynamics of sensilla evolution as pheromone blends diverge.

## Acknowledgments

We would like to thank Lynn Faust for collection material from the Great Smokey Mountains, and Rebecca Clement and Samantha Standring for collection material from Cameroon. Thank you to Eric Fromm and the Georgia Electron Microscopy Laboratory for SEM imaging training and troubleshooting. Thank you to Jim Leebens-Mack, Sarah E. Lower, and Shu-Mei Chang for their comments on an early version of this manuscript.

Mention of trade names or commercial products in this publication is solely for the purpose of providing specific information and does not imply recommendation or endorsement by the USDA; USDA is an equal opportunity provider and employer.

## Data Availability

The data underlying the results presented in this study are available from Dryad.

## Supporting information

**Figure S1. Diurnal species antennal area.** (A) In diurnal species, males have significantly larger antennal areas than females (x̵_Male_=1.812± 0.66 mm^2^, x̵_Female_=1.2±0.38 mm^2^; Student’s t ratio = -3.86, p=0.0005).

**Figure S2: Total sensilla densities by activity time and sex.** (A) Sensilla densities by activity time: there is no significant difference in total sensilla density between diurnal and nocturnal species (x̵_Diurnal_ ==3392±907 per mm^2^, x̵_Nocturnal_ =2949±1766 per mm^2^; DFDen=5, F=0.187, p=0.683); (B) Sensilla densities by sex: there is no significant difference in total sensilla density between females and males (x̵_Male_=3414±1720 per mm^2^, x̵_Female_=2864±1128 per mm^2^; DFDen=38, F= 2.92, p=0.0956); (C). Sensilla densities between nocturnal females and males: nocturnal males have significantly higher density of sensilla than nocturnal females (x̵_Male_=3503±2145 per mm^2^, x̵_Female_=2395±1118 per mm^2^; Student’s t ratio= -3.17, p=0.003).

**Figure S3.. Chemosensilla counts by sex and activity** (A) Chemosensilla counts of diurnal species by sex: diurnal males have significantly more chemosensilla than diurnal females (x̵_DMale_ = 1862±1326, x̵_DFemale_ = 660±259; Student’s t-ratio= -4.57, p<0.0001); (B) Chemosensilla counts of males by activity time: diurnal males have significantly more chemosensilla than nocturnal males: (x̵_DMale_ = 1862±1326, x̵_NMale_ = 682±195; Student’s t-ratio= 3.08, p=0.0155); (C) Chemosensilla counts of females by activity time: there is no significant difference in chemosensilla counts between nocturnal and diurnal females (x̵_DFemale_ = 660±259, x̵_NFemale_= 507±334; Student’s t-ratio= 0.4, p=0.7); (D) Chemosensilla counts of nocturnal species by sex: there is no significant difference in sensilla counts between nocturnal females and males (x̵_NMale_= 682±195, x̵_NFemale_= 507±334; Student’s t-ratio= -0.77, p=0.45).

**Figure S4. Antennal area and mechano- and chemosensilla counts and densities.** (A) Mechanosensilla counts are positively correlated with antennal area (R^2^=0.313, p=0.0001). (B) Chemosensilla counts are positively correlated with antennal area (R^2^=0.223, p=0.0016). (C) Mechanosensilla density is negatively correlated with antennal area (R^2^= 0.549, p=0.0001). (D) Chemosensilla density is negatively correlated with antennal area (R^2^= 0.199, p=0.0031).

**Figure S5. Mechanosensilla densities by sex and activity.** (A) Mechanosensilla density of diurnal species by sex: diurnal females have a significantly greater density of mechanosensilla than diurnal males (x̵_DFemale_= 2887±571 per mm^2^, x̵_DMale_= 2265±954 per mm^2^; Student’s t-ratio=2.7, p=0.0109); (B) Mechanosensilla density of nocturnal species by sex: nocturnal males have a greater density of mechanosensilla than nocturnal females (x̵_NFemale_ = 1870±896 per mm^2^, x̵_NMale_ = 2265±1243 per mm^2^; this trend was marginally significant: Student’s t-ratio= -1.97, p=0.057); (C) Mechanosensilla density of females by activity time: there is no significant difference in mechanosensilla densities between diurnal and nocturnal females (x̵_DFemale_= 2887 ±571per mm^2^, x̵_NFemale_=1871±896per mm^2^; Student’s t-ratio=1.35, p=0.0.23); (D) Mechanosensilla density of males by activity time: there is no significant difference in mechanosensilla density between diurnal and nocturnal males (x̵_DMale_=2265± 954 per mm^2^, x̵_NMale_ =2265±1243 per mm^2^; Student’s t-ratio= -0.00, p=0.999).

**Figure S6. Distribution of mechanosensilla across the antenna.** Average sensilla counts per antennomere for females (red) and males (black) for each species. (*L.* = *Lucidota*, *P.* = *Photinus*, *Py.* = *Pyropyga*, *Ph. = Photuris*).

**Figure S7: Distribution of chemosensilla across the antenna.** Average sensilla counts per antennomere for females (red) and males (black) for each species (*L.* = *Lucidota*, *P.* = *Photinus*, *Py.* = *Pyropyga*, *Ph. = Photuris*).

**Figure S8. Distribution of unique sensilla of *Lucidota* across the antenna.** Average sensilla counts per antennomere for females (red) and males (black) (*L.* = *Lucidota*).

**Table S1. Voucher specimens used for SEM imaging.** Species name, specimen number, KSH voucher ID number, and sex.

**Table S2. Mechanosensilla counts.** Individual mechanosensilla (C1, C2, SC) counts (mean ± stdev) for each species (F: 3 females, M: 3 males, D: diurnal, N: Nocturnal, *L.* = *Lucidota*, *P.* = *Photinus*, *Py.* = *Pyropyga*, *Ph. = Photuris*).

**Table S3. Mechanosensilla densities.** Individual mechanosensilla (C1, C2, SC) density (mean ± stdev N/mm^2^) of each species (F: 3 females, M: 3 males, D: diurnal, N: Nocturnal, *L.* = *Lucidota*, *P.* = *Photinus*, *Py.* = *Pyropyga*, *Ph. = Photuris*). (-) type absent.

**Table S4. Chemosensilla counts.** Individual chemosensilla (B1-B3, B7, B10-B13, T1) counts (mean ± stdev) for each species (F: 3 females, M: 3 males, D: diurnal, N: Nocturnal, *L.* = *Lucidota*, *P.* = *Photinus*, *Py.* = *Pyropyga*, *Ph. = Photuris*).

**Table S5. Chemosensilla densities.** Individual chemoreceptor (B1-B3, B7, B10-B13, T1) density (mean ± stdev) of each species (F: 3 females, M: 3 males, D: diurnal, N: nocturnal, *L.* = *Lucidota*, *P.* = *Photinus*, *Py.* = *Pyropyga*, *Ph. = Photuris*). (-) type absent.

**Table S6. X1 sensilla counts and densities.** Mean counts and densities (mean ± standard deviation) for X1 sensilla by species and sex. *X1 sensilla were not present in all specimens within a species, specimens lacking these sensilla were excluded from mean and standard deviation calculations. (F: 3 females, M: 3 males, D: diurnal, N: nocturnal, *L.* = *Lucidota*, *P.* = *Photinus*, *Py.* = *Pyropyga*, *Ph. = Photuris*). (-) type absent.

